# MORC2 phosphorylation fine tunes its DNA compaction activity

**DOI:** 10.1101/2024.06.27.600912

**Authors:** Winnie Tan, Jeong Veen Park, Hariprasad Venugopal, Jie Qiong Lou, Prabavi Shayana Dias, Pedro L. Baldoni, Toby Dite, Kyoung-Wook Moon, Christine R. Keenan, Alexandra D. Gurzau, Andrew Leis, Jumana Yousef, Vineet Vaibhav, Laura F. Dagley, Ching-Seng Ang, Laura Corso, Chen Davidovich, Stephin J. Vervoort, Gordon K. Smyth, Marnie E. Blewitt, Rhys S. Allan, Elizabeth Hinde, Sheena D’Arcy, Je-Kyung Ryu, Shabih Shakeel

## Abstract

Variants in the poorly characterised oncoprotein, MORC2, a chromatin remodelling ATPase, lead to defects in epigenetic regulation and DNA damage response. The C-terminal domain (CTD) of MORC2, frequently phosphorylated in DNA damage, promotes cancer progression, but its role in chromatin remodelling remains unclear. Here, we report a molecular characterisation of full-length, phosphorylated MORC2, demonstrating its preference for binding open chromatin and functioning as a DNA sliding clamp. We identified a phosphate interacting motif within the CTD that dictates ATP hydrolysis rate and cooperative DNA binding. The DNA binding impacts several structural domains within the ATPase region. We provide the first visual proof that MORC2 induces chromatin remodelling through ATP hydrolysis-dependent DNA compaction, regulated by its phosphorylation state. These findings highlight phosphorylation of MORC2 CTD as a key modulator of chromatin remodelling, presenting it as a potential therapeutic target.

## Introduction

The human Microrchidia CW-type zinc finger (MORC) is a family of four nuclear proteins (MORC1-4) that maintain chromatin structure and function^1–3^. All MORCs share three unifying features: an N-terminal **g**yrase, **h**eat shock protein 90, histidine **k**inase and Mut**L** (GHKL) ATPase domain, a central CW-type zinc finger (CW) domain, and a divergent C-terminal region with predicted coiled coils (CCs)^4^. While the GHKL and CW domains are predicted to be important for transcriptional silencing and chromatin binding respectively^5,6^, the CC domain is proposed to facilitate MORC’s homo-dimerisation^4^. Crystal structures of ATPase domains of MORC2, MORC3 and MORC4 reveal their propensity to dimerise upon ATP binding and/or DNA interaction^2,3,7^, but do not explain whether this dimerisation is sufficient to drive chromatin remodelling. This is partly due to a lack of structures of full-length MORCs, which are difficult to produce recombinantly because of the large, disordered C-terminal domain (CTD). Notably, extensive phosphorylation of MORC2 in its CTD (**Fig 1a**) suggests regulation of its functions via phosphorylation^8^. Phosphorylated MORC2 is associated with DNA damage sites^9^ and phosphorylation of S739 by PAK1 kinase is implicated in gastric cancer progression^8^. Moreover, MORC2 overexpression has been linked to breast and liver cancers, although the role of MORC2 phosphorylation in these contexts is unknown^8,10,11^.

**Figure 1.**
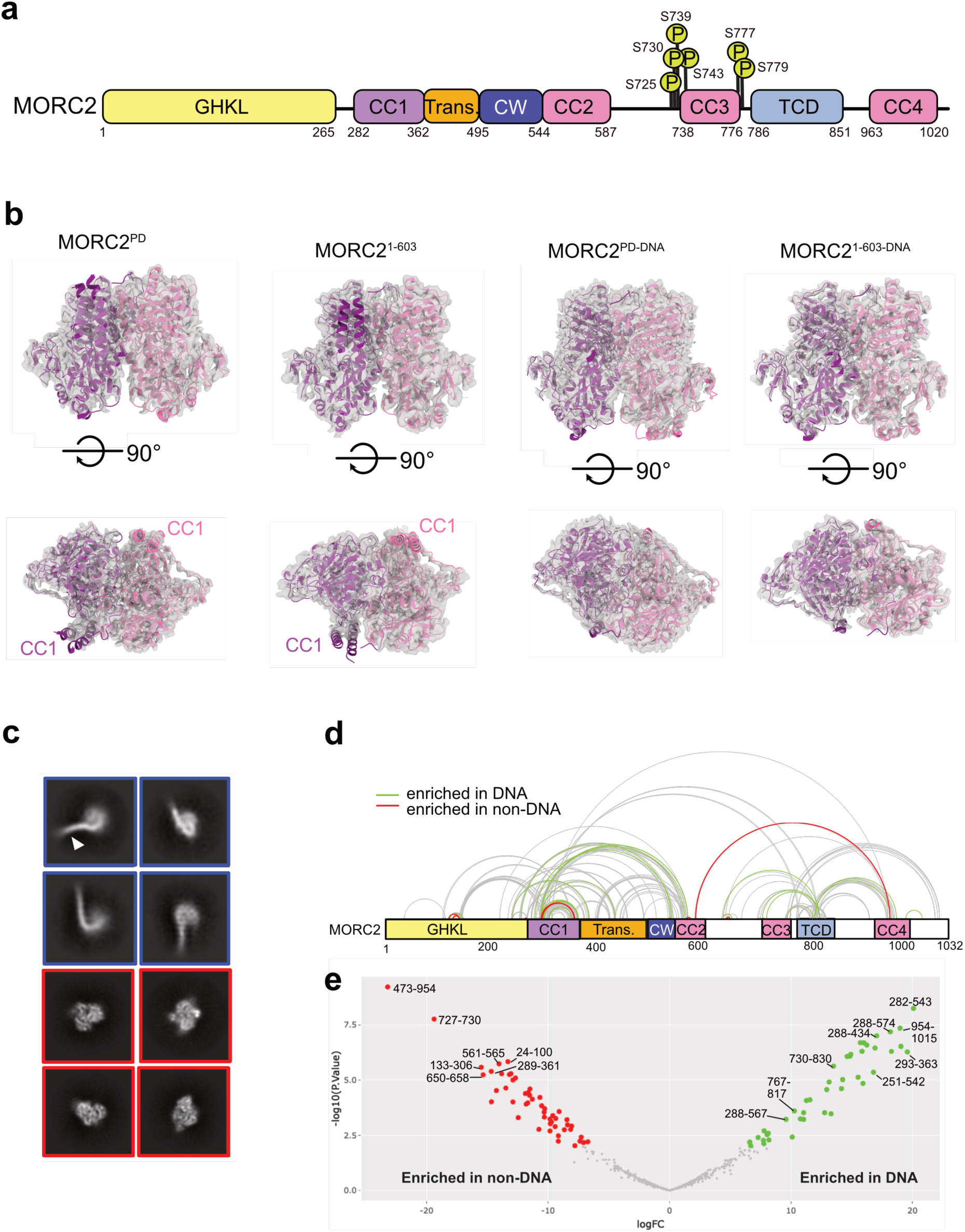
CryoEM structures of human MORC2. **a.** Domain diagram of MORC2. The GHKL indicates Gyrase, Hsp90, Histidine Kinase, MutL domain, CC1-4 indicates coiled-coil domains, Trans indicates Transducer-like domain, CW indicates CW-type zinc finger domain and TCD indicates Tudor-chromodomain. Phosphorylation sites studied in this work are marked as ‘P’ with the residue number indicated. **b.** The cryoEM structures of phosphodead (MORC2^PD^) mutant, MORC2 ATPase (MORC2^1-603^), phosphodead MORC2 with DNA (MORC2^PD-DNA^), and MORC2 ATPase with DNA (MORC2^1-603-DNA^). Coiled-coil 1 is marked as CC1, which is only seen in DNA-free samples. Pink and purple colour models fitted in the cryoEM maps indicate one protomer each of the MORC2 homodimer. **c**. Selected 2D reference-free class averages of MORC2^PD-DNA^. The blue box shows the 2D classes where DNA (marked by white arrowhead in one of the classes) is seen bound to fuzzy MORC2 density. The red box indicates high-resolution 2D classes where secondary structure elements are clearly visible. **d.** Map of quantified cross-links in MS analysis of MORC2 DNA and non-DNA bound samples. Crosslinks with a similar abundance in DNA and non-DNA (grey lines), cross-links considered significantly enriched (adjusted *p*-value ≤ 0.05) in DNA (green lines) and in non-DNA (red lines) are shown. **e.** Volcano plot of quantified cross-links, where the log2 DNA/non-DNA fold changes are plotted against the -log10 p-value. Crosslinks that were considered significantly enriched (adjusted *p*-value ≤ 0.05) in DNA (green) and in non-DNA (red) are highlighted; some of the top hits have the peptide residue numbers indicated. The full list of all enriched crosslinks is in Supplementary Data Table 3.

## Results

### Recombinant MORC2 is phosphorylated at multiple sites

To understand the molecular mechanism of MORCs and how they are regulated, we purified recombinant, functionally active, full-length MORC2 from insect cells (**Extended Data Fig 1a**) and conducted a comprehensive analysis employing structural, biochemical, biophysical, single molecule, and cellular approaches. Size exclusion chromatography multi-angle light scattering (SEC-MALS) analysis of full-length MORC2 reveals a homodimer of 264 kDa, which differs from the ATPase domain (1-603 amino acids) that dimerises only when ATP analog, AMP-PNP, is present (**Extended Data Fig 1b**). SEC-MALS also showed that all MORC2 truncation variants that contained CTD formed dimers (**Extended Data Fig 1c**), confirming that MORC2 CTD enables its dimerisation. To observe whether MORC2 forms dimers in cells, we used Number and Brightness (NB) measurement of the eGFP tagged MORC2 plasmid in a MORC2 knockout HEK293T cell line. About 20% of the MORC2 population is present as dimer, and this doubled to ∼40% upon addition of neocarzinostatin, a DNA damage agent that is known to cause double stranded (ds) DNA breaks (**Extended Data Fig 1d-f**). These data suggest dimerised MORC2 may be the functional form, at least in DNA repair.

The purified MORC2 came phosphorylated from insect cells, and the level of phosphorylation further increased upon incubation with PAK1 kinase^9^ (**Supplementary Data Table 1**). Tandem mass spectrometry (MS/MS) analysis revealed extensive phosphorylation at multiple sites within MORC2, with the top 6 hits S725, S730, S739, S743, S777, and S779, which were further phosphorylated by the PAK1 kinase. Of these residues, all except S725 are evolutionarily conserved and reside within the CTD region, which we coined as phosphate interacting motif (PIM) (**Extended Data Fig 2a, Extended Data Fig 3**). To evaluate the impact of phosphorylation on MORC2’s DNA binding capability, we incubated the purified MORC2 with λ-protein phosphatase (λPPase) before performing an electrophoretic mobility shift assay (EMSA) in the presence of double-stranded DNA (dsDNA). EMSA showed increased retention of dephosphorylated MORC2 on DNA compared to the wild-type and PAK1-treated MORC2 (**Extended Data Fig 2b-c**) suggesting dephosphorylated MORC2 may be a stronger DNA binder.

### MORC2 undergoes subtle structural changes upon DNA binding

To determine how phosphorylation of MORC2 affects DNA binding, we purified a MORC2 phosphodead (MORC2^PD^) mutant comprising S725A, S730A, S739A, S743A, S777A and S779A mutations (**Fig 1a, Extended Data Fig 1a**). We determined the structures of MORC2^PD^ and MORC2 ATPase domain (MORC2^1–603^) in the presence of AMP-PNP (MORC2^PD^, MORC2^1–603^) and with 60 base pairs (bp) dsDNA (MORC2^PD-DNA^, MORC2^1–603-DNA^) at overall resolutions of 2.4, 3.2, 1.9 and 2.5 Å, respectively, using cryo-electron microscopy (cryoEM) (**Fig 1b, Extended Data Fig 4a-b, Supplementary Data Table 2**). The dimerised ATPase domain is well-resolved in all the cryoEM maps, but no density is seen for CTD in the full-length MORC2^PD^ and MORC2^PD-DNA^. We also determined a 2.5 Å cryoEM structure of full-length MORC2 ATPase-dead mutant (MORC2^S87A^) in the presence of AMP-PNP, where like other cryoEM maps only the ATPase domain was resolved (**Extended Data Fig 5a-d**). The AMP-PNP density near the ATP lid was visible in MORC2^S87A^ map confirming that this mutant can bind but not hydrolyse ATP, mimicking S87L variant where some residual ATPase activity is still detected (**Extended Data Fig 5e**). The full-length WT MORC2 displayed a preferred orientation, and we could not resolve it beyond 2D classification (**Extended Data Fig 6a-b**). In the MORC2^PD^ and MORC2^1–603^ constructs, we observed a coiled-coil region (CC1) that was absent in their DNA-bound states (**Fig 1b**, **Extended Data Fig 4c**). We used crystal structures of MORC2^1–603^ protein (PDB:5OF9 and 5OFB)^7^ to build models into our cryoEM maps, which showed no noticeable difference between the two. The MORC2^PD-DNA^ showed some 2D classes where it was bound to the DNA (**Fig 1c**), but as for full-length MORC2, we could not resolve these 2D classes into a 3D map due to low resolution and absence of diverse views (**Extended Data Fig 6a-b)**. Interestingly, almost all CC1 was missing in the DNA-bound cryoEM structures indicating high flexibility of this region upon DNA binding. Overall, these structures show that the core of the ATPase domain does not undergo large conformational changes in the presence of DNA, and that MORC2 CTD is flexible.

Since cryoEM did not resolve the MORC2 structure beyond the N-terminal ATPase domain, we employed quantitative crosslinking mass spectrometry to investigate conformational changes in MORC2 induced by DNA binding. We used a high density crosslinker, sulfosuccinimidyl-4,4′-azipentanoate (‘sulfo-SDA’), where its NHS ester group binds to primary amines and the diazarine ring to the side chain of any residue within its vicinity. We crosslinked full-length MORC2 on its own (MORC2^Apo^), in the presence of AMP-PNP with DNA (MORC2^AMPPNP-DNA^) and without DNA (MORC2^AMPPNP^).

There were 337 crosslinks in the MORC2^Apo^, 214 in MORC2 ^AMPPNP^, and 359 in MORC2^AMPPNP-DNA^, at 5% False Discovery Rate (FDR, **Supplementary Data Table 3**). We used the crosslinks from MORC2^AMPPNP-DNA^ to validate the AlphaFold2 multimer-predicted models for full-length MORC2 dimer. Model 1 closely resembles the MORC2 crystal structure (PDB:5OF9) and was used for validation by crosslinking mass spectrometry data (**Extended Data Fig 6c-e**). We used XMAS^12^, which identifies homodimeric interaction based on peptide sequence overlap, to map the crosslinks on Model 1 within the proximity of 20 Å (**Extended Data Fig 6f**). There were 64 inter-molecular crosslinks identified between the two protomers. There were 7 crosslinks between the CTD (residues 904-1029) from the two protomers, which are paired next to each other in the model, thus confirming that they are coiled coils (CC4). A large cluster of crosslinks was observed within the ATPase domain, indicating that it is the most well folded domain across all the regions of MORC2, consistent with our cryoEM structures where we were able to resolve the ATPase domain, with the rest being too flexible.

We quantified the crosslinks for a statistical-based comparison of MORC2^Apo^, MORC2^AMPPNP^ and MORC2^AMPPNP-DNA^ conformational changes (**Supplementary Data Table 3**). Between MORC2^Apo^ and MORC2^AMPPNP^, there were 93 common crosslinks, 30 unique crosslinks in MORC2^Apo^ and 47 in MORC2^AMPPNP^. These unique crosslinks appeared throughout the length of MORC2, indicating that ATP dimerisation leads to conformational changes in other regions of the protein (**Extended Data Fig 7a-b**). Without ATPase dimerisation, the CC1 domain interacts with the CC3 domain (**Extended Data Fig 7a**). However, upon dimerisation, CC1 contacts the TCD domain, as evidenced by enrichment of crosslinks in the AMP-PNP sample (**Extended Data Fig 7a, Supplementary Data Table 3**). The most striking domain-level changes involved crosslinks in the GHKL domain near S87, ATP binding site where we observed 32 crosslinks in the GHKL domain in presence of AMP-PNP (**Extended Data Fig 7c, Supplementary Data Table 3**).

The MORC2^AMPPNP^ and MORC2^AMPPNP-DNA^ share many crosslinks suggesting that the two complexes adopt similar conformations. However, by focusing on significantly enriched crosslinks between MORC2^AMPPNP^ and MORC2^AMPPNP-DNA^, the abundance of some crosslinks differed, suggesting subtle changes in MORC2 conformation upon DNA binding (**Fig 1d-e**). In particular, the residues within CC1 appear to make different contacts between the two samples. Also, MORC2^AMPPNP^ has long range contacts including 176-749, 290-561 and 473-954, which disappear in presence of DNA (**Fig 1d**). By mapping the crosslinks on the MORC2 Model 1, we showed that the CC1 domain crosslinked to the CW-CC2 domains upon DNA binding (**Extended Data Fig 6f, Supplementary Video 1**), indicating that MORC2 CC1 engaged with a patch on the CW-CC2 domain that was distal to the dimerisation interface mediated by AMP-PNP. Enrichment of these crosslinks suggests that the CC1 region is more accessible in the absence of DNA.

### MORC2 has multiple DNA binding sites

To further characterise MORC2 DNA binding, we performed hydrogen-deuterium exchange mass spectrometry (HDX) on MORC2^1–603^ and MORC2 CTD (MORC2^496–1032^) with and without 60 bp dsDNA in the presence of AMP-PNP (**Supplementary Data Table 4**). In MORC2^1–603^, we observed a significant increase in deuterium uptake in the ATP lid upon DNA binding (**Fig 2a**, peptides 81 to 109). Our MORC2^1–603^ cryoEM structure and the previous crystal structure (PDB:5OF9)^15^ showed that upon AMP-PNP binding, the ATP lid covers the active site and modulates rotation of CC1, which contributes to the MORC2 dimerisation interface. Our ATPase-dead S87A mutant (**Extended Data Fig 5b**) and S87L Charcot-Marie-Tooth patient mutant^15^ structures also show a change in the ATP lid. Thus, the DNA-induced, altered conformation of the ATP lid detect by HDX could have a functional consequence for MORC2 chromatin remodelling.

**Figure 2.**
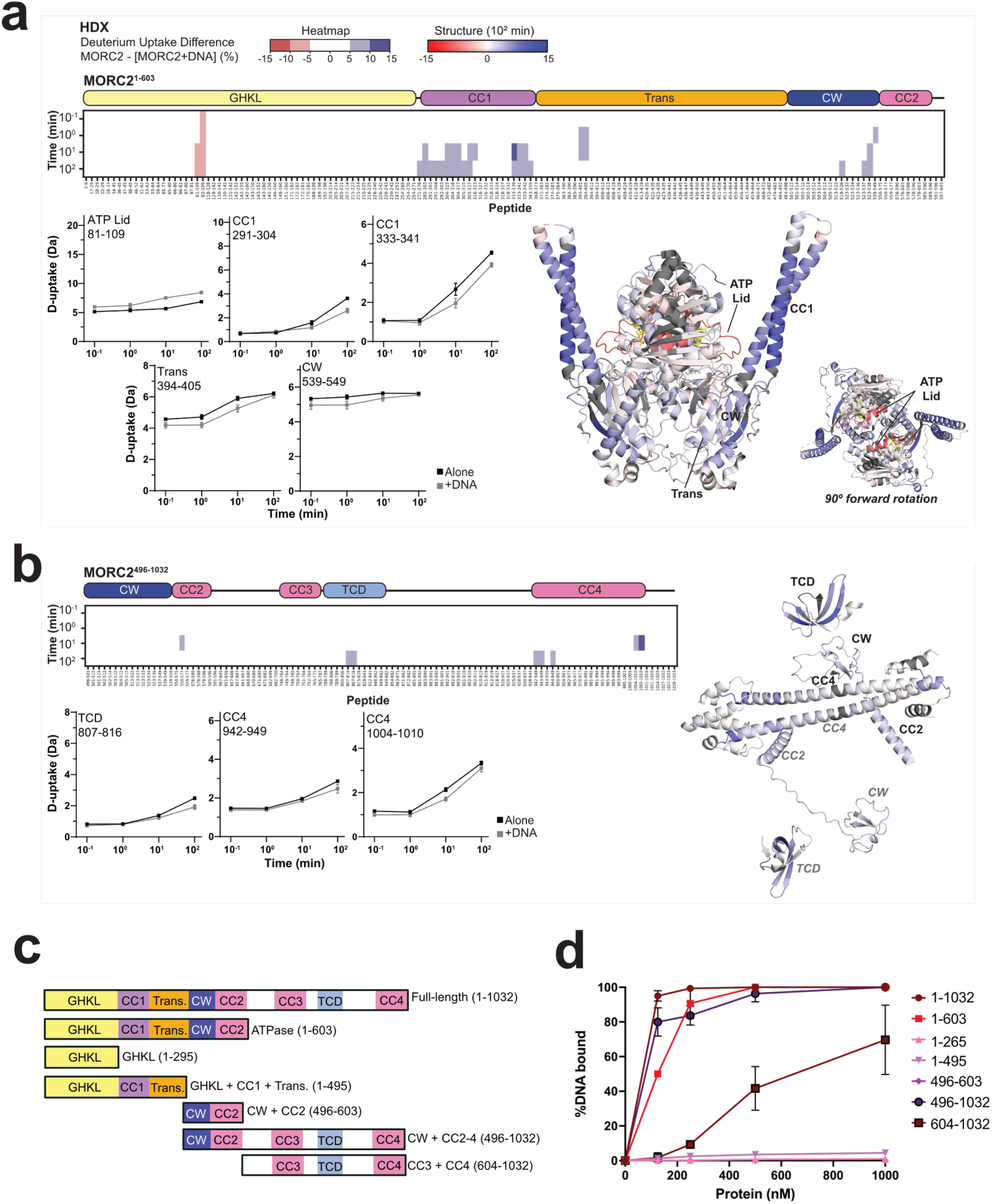
DNA binding by MORC2 ATPase and CTD domains in solutions. **a.** Heatmap showing the difference in deuterium uptake between MORC2^1-603^ alone and with three-fold excess 60 bp dsDNA (top). Deuterium uptake plots for example peptides (left). The MORC2 ATPase crystal structure (PDB: 5OF9) coloured by difference after 10^2^ min exchange based on DynamX residue-level scripts without statistical filters (right). Residues without coverage are grey. **b.** Heatmap showing the difference in deuterium uptake between MORC2^496–1032^ alone and with three-fold excess 60 bp dsDNA (top). Deuterium uptake plots for example peptides (left). The MORC2^496–1032^ Alphafold Model 1 coloured by difference after 10^2^ min exchange based on DynamX residue-level scripts without statistical filters (left). Residues without coverage are grey. For the model, only residues with >40 confidence are shown. For **a** and **b**, blue/red heatmap colouring indicates a difference ≥5% with a *p*-value ≤0.01 in Welch’s t-test (n=3). Plots show deuterium uptake for representative peptides for MORC2 alone (black) or with DNA (grey). Error bars are ±2 standard deviation (SD, n=3 or 4) and the y-axis is 80% of the maximum theoretical deuterium uptake, assuming the complete back exchange of the N-terminal residue. **c.** Schematic for MORC2 constructs used in DNA binding EMSA. **d.** Quantification of percentage of 60bp dsDNA bound to MORC2 FL (1-1032), ATPase (1-603), GHKL (1-265), GHKL+CC1 (1-495), CW domain (496-603), CW+CC1+CTD (496-1032) and CCW+CTD (604-1032). The points are shown as mean +/- standard deviation (n=3).

The presence of 60 bp dsDNA with MORC^1–603^ also reduced deuterium uptake in CC1 and nearby regions, transducer-like and CW. This suggests CC1 and nearby regions form a high-affinity DNA binding site, somewhat consistent with previous mutagenesis of the CC1 loop (point mutation of R326, R329 and R333) that prevented DNA binding^15^. We saw only modest decreases in deuterium uptake in the CC1 loop (below our significance thresholds) as this loop is disordered and reached maximal deuteration at our initial timepoint. The reduced deuterium uptake in CC1 and CW may also be attributed to an intra-molecular interaction induced by DNA, consistent with our crosslinking experiments (**Fig 1d-e**). Notably, we did not see a DNA-induced change in deuterium uptake in the CW when CC1 was not present in MORC2^496–1032^. For MORC2^496–1032^, we saw reduced deuterium uptake in the TCD and at opposite ends of CC4 (**Fig 2b**), hinting that the CTD contains secondary, low-affinity DNA binding sites.

To further investigate our HDX observations, we performed EMSA with various MORC2 truncations, in the presence of 60 bp dsDNA (**Fig 2c**). All constructs that lacked CW and CC2 showed lower DNA binding in comparison to the full-length MORC2 (**Fig 2d, Extended Data Fig 8a**). The constructs containing only GHKL and CC1 domains, or only CW and CC2, showed negligible DNA binding, whereas some DNA binding was rescued in the 604-1032 construct that lacks CC1-2. Taken together, the HDX and EMSA data show that CC1 DNA binding requires CW-CC2, and that CC1 and CW-CC2 alone are insufficient for DNA binding. We further tested the DNA binding ability of isolated CC2, CC3 and CC4. Interestingly, CC2 (544-587), and CC4 (963-1020) domains were able to bind DNA but not the CC3 region that contains the PIM domain (**Extended Data Fig 8b-c**). This is consistent with the HDX data where the CW adjacent to CC2 has measurable DNA binding (**Fig 2b**). Thus, this work confirms that MORC2 has multiple DNA binding sites in the ATPase and CTD domains.

### MORC2 ATPase activity is phosphorylation-dependent

The rate of ATP hydrolysis by chromatin remodellers may have an impact on chromatin regulation^2,3^. As the MORC2 proteins purified from insect cells were phosphorylated, we hypothesised that MORC2 phosphorylation may affect its function. We performed Fluorescence Polarisation ATPase assays on MORC2^WT^, MORC2^1–603^ and MORC2^PD^. MORC2^WT^ and MORC2^1–603^ were able to hydrolyse ATP at a turnover rate (*k*_cat_) of 0.06 ± 0.02 µM ADP/min/µM and 0.05 ± 0.02 µM ADP/min/µM, which is similar to other GHKL ATPases^2,3,13^. Surprisingly, MORC2^PD^ hydrolysed ATP ∼2-fold faster than MORC2^WT^ at *k*_cat_ of 0.11 ± 0.03 µM ADP/min/µM (**Fig 3a**), and addition of DNA did not alter MORC2 ATPase activity (**Extended Data Fig 8d-e**). To assess the DNA binding activity of MORC2, we performed surface plasmon resonance on MORC2^WT^, MORC2^1–603^ and MORC2^PD^ mutant. We found all MORC2 variants were able to bind to 60 bp DNA within a similar nanomolar affinity range (**Fig 3b**). Together, these data show that the MORC2 CTD, which harbours the phosphorylation sites, regulates the ATP hydrolysis activity of MORC2 independently of DNA binding.

**Figure 3.**
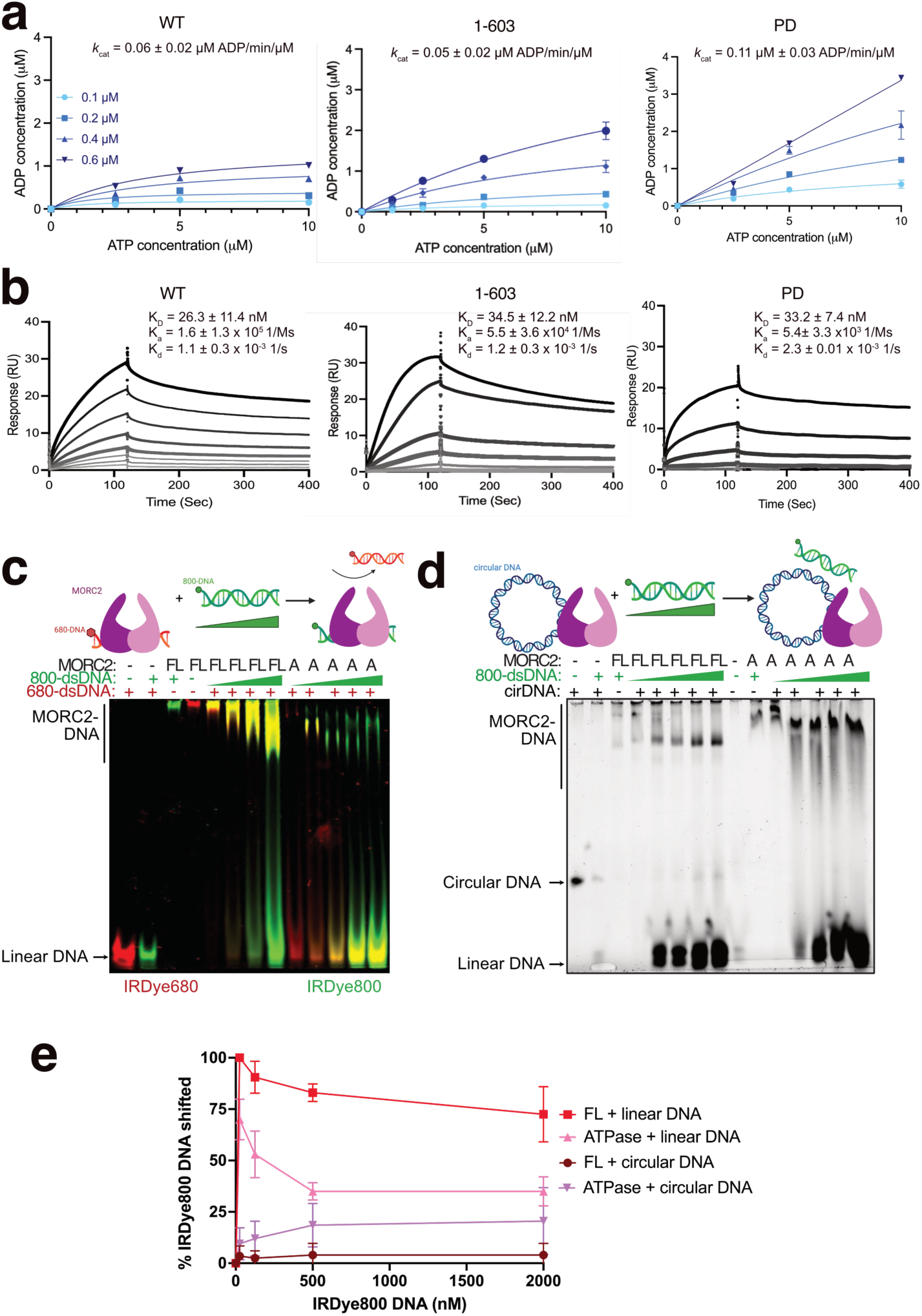
MORC2 interacts with DNA at multiple binding sites independent of ATP binding. **a.** *In vitro* Fluorescence Polarisation ATPase activity of MORC2 WT, 1-603 and PD mutant. Individual measurements (n=3) are shown as points, and the solid line represents the non-linear fit of the data. The indicated k_cat_ values are mean ± standard error of mean (SEM) representative of three experiments. **b.** Surface plasmon resonance analysis of MORC2 WT, ATPase (1-603) and PD mutant with protein concentrations of 0, 4, 8, 16, 33, 63, 125, 250 and 500 nM. Values for K_D_ (equilibrium dissociation constant), K_a_ (association rate constant) and K_d_ (dissociation rate constant) are shown as mean ± SEM, representative of three experiments. **c.** A 60-bp linear IRDye680-labelled dsDNA was used to pre-incubate with 100 nM MORC2 full-length (FL) or ATPase (A) construct, followed by 0, 25, 125, 500 and 2000 nM of 60-bp IRDye800-labelled linear dsDNA. Overlays of IRDye680 (red) and IRDye800 (green) channel images are shown. **d.** A 101-bp circular dsDNA was used to incubate 100 nM MORC2 full-length (FL) or ATPase (A) construct, followed by 0, 25, 125, 500 and 2000 nM IRDye-800 linear dsDNA. SYBR gold staining of 6% PAGE gel is shown. **e.** Quantification of IRDye-800 DNA shifted for the linear and circular DNA binding competition assay. The points are shown as mean ± SD (n=2).

### Full-length MORC2 acts as a DNA sliding clamp

To understand MORC2 interaction with DNA, we first determined the minimal length of DNA required for binding. We performed EMSA with MORC2^WT^ and MORC2^1–603^ in the presence of IRDye-700 labelled 10, 14, 19, 24, 29, 34, 39, 44 and 60 bp dsDNA. Both constructs of MORC2 bind to dsDNA longer than ∼29 bp (**Extended Data Fig 9**). Next, we investigated whether MORC2 locks onto DNA or slides off as proposed for MORC3^14^. We first assembled MORC2^WT^ or MORC2^1–603^ on 60 bp linear or unlabelled 101 bp circular plasmid DNA for 30 minutes at room temperature, then added 20-fold molar excess of IRDye800-labelled 60 bp dsDNA and monitored the DNA binding after 10 minutes. About 74% of the competitive DNA is incorporated into the linear DNA-MORC2^WT^ complex with only 4% in the circular DNA complex (**Fig 3c-d**). In comparison, 50% of the linear DNA-MORC2^1–603^ complex had shifted compared to 25% of the circular DNA complex in presence of competitive DNA (**Fig 3e, Extended Data Fig 10a-b**). This shows that the additional DNA binding sites on CTD detected by HDX are responsible for formation of a tight clamp. Additionally, these results suggests that the clamp dissociates from linear DNA by sliding off the end and not by opening up because in the latter case it would dissociate from circular DNA as well, hence, indicating that MORC2 is likely a DNA sliding clamp.

### MORC2 is predominantly associated with accessible chromatin

To identify the preferred genomic regions of MORC2 binding and its impact on chromatinisation, we performed chromatin immunoprecipitation sequencing (ChIP-seq) in the HEK293T cell line utilising antibodies specific for MORC2 and H3K9me3, a mark of constitutive heterochromatin, since MORC2 is known to associate with the human silencing hub complex^15^. MORC2 binding sites have previously been reported to be enriched at promoters in the HeLa cell line^15^. Therefore, we also performed assays for transposase-accessible chromatin with sequencing (ATAC-seq) to determine if MORC2 was binding open chromatin. We identified 15,366 MORC2 binding sites in HEK293T cells of which 8,317 overlapped gene promoters according to ChIP-seq (**Fig 4a**). The majority of such regions were accessible in ATAC-seq and depleted of H3K9me3 (**Fig 4b-c**). In contrast, 3,095 MORC2 peaks overlapped H3K9me3 outside of promoter regions, indicating binding in heterochromatin (**Extended Data Fig 10c-d**) and hinting a weaker colocalization between MORC2 and H3K9me3 regions. To further explore the global binding profile of MORC2, we examined the relationship of MORC2 binding sites with other genomic locations such as repetitive elements including LINE-1 elements and endogenous retroviruses (ERV) where MORC2 is also known to play a role^15^; however we found limited binding at these repetitive elements (**Fig 4d)**. A similar observation was reported previously, where out of 4,500 MORC2 peaks, only 490 peaks overlapped with H3K9me3^15^, suggesting a non-canonical epigenetic silencing pathway mediated by MORC2.

**Figure 4.**
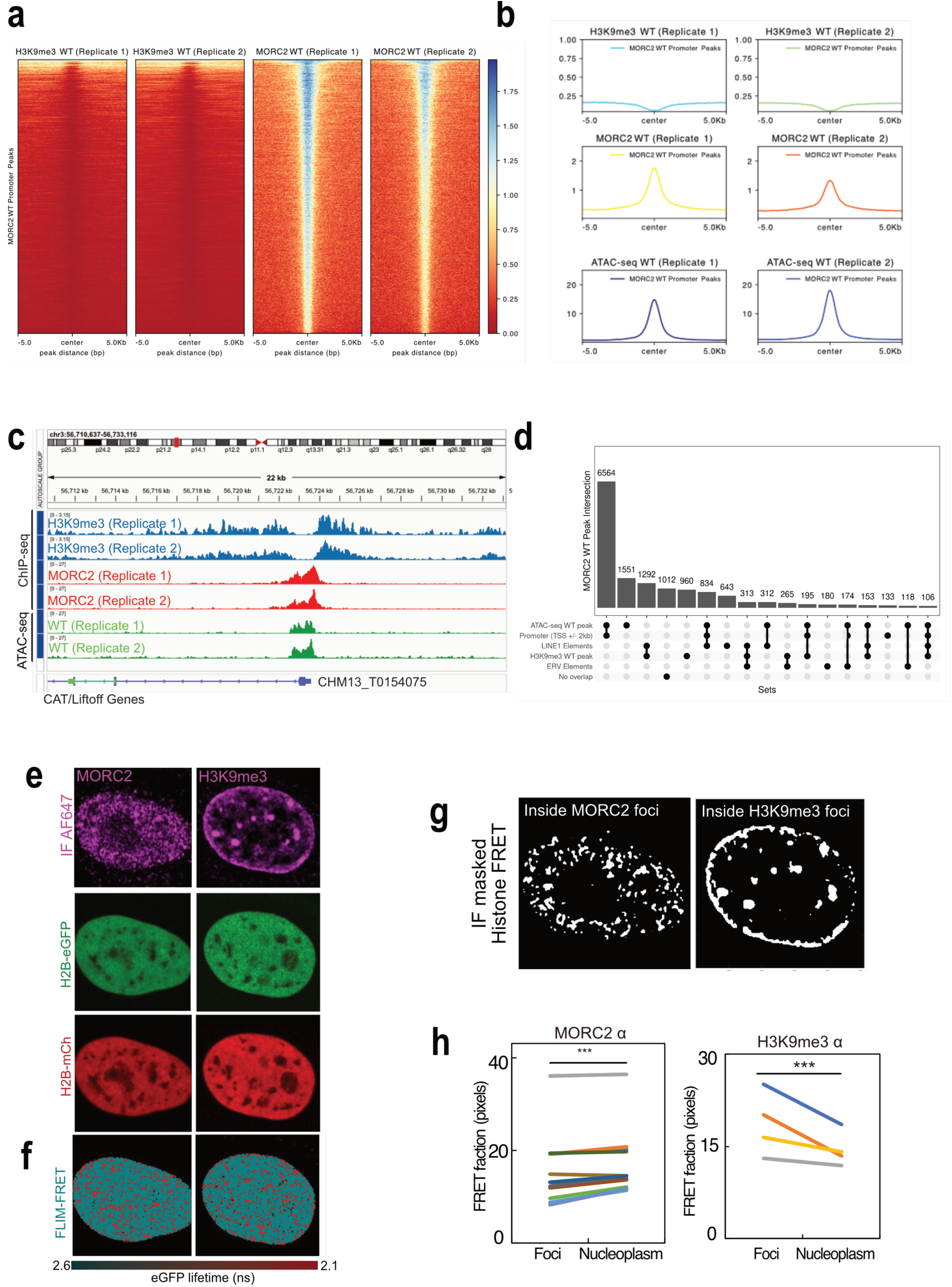
MORC2 associates with open chromatin region. **a.** Heatmaps showing H3K9me3 and MORC2 ChIP-seq coverage of promoter-overlapping MORC2 WT peaks (total number of peaks = 8317), sorted by MORC2 intensity. **b.** Mean coverage of H3K9me3 and MORC2 ChIP-seq and ATAC-seq signals from 5kb upstream of promoter-overlapping MORC2 WT peaks (total number of peaks = 8317) to 5 kb downstream. **c.** Representative Integrative Genomics Viewer (IGV) image of H3K9me3 and MORC2 ChIP-seq and ATAC-seq tracks in HEK293T cells showing chr3:56,710,637-56,733,116. For (**a-c**), two biological replicates from ChIP- and ATAC-seq assays were used for visualization purposes. **d.** Upset plot of MORC2 WT peaks showing the number of peaks overlapping H3K9me3 peaks, promoter regions and ATAC-seq peaks. The number of peaks without any overlap is also presented. **e.** FLIM-FRET analysis of HEK293T co-expressing H2B-eGFP and H2B-mCherry, with MORC2 and H3K9me3 immunofluorescence labelled with Alexa Fluor dye 647 (AF647). **f.** The corresponding pseudo-coloured maps show chromatin compaction, where red colour represents compact chromatin region. **g.** Masks based on MORC2 or H32K9me3 intensity image mapped to H2B FRET map. **h.** Quantification of the fraction of histone FRET (compact chromatin) inside versus outside of MORC2 or H3K9me3 foci across multiple cells (N = 20 cells for MORC2 and N = 4 cells for H3K9me3, two biological replicates). Foci represent compact chromatin and nucleoplasm represents open chromatin; ns indicates *P*> 0.05, *** indicates *P*<0.001, and **** indicates *P*<0.0001, paired *t*-test.

To directly visualise MORC2 binding to chromatin, we used fluorescence lifetime imaging microscopy (FLIM)-coupled Förster resonance energy transfer (FRET) between fluorescently labelled histones (eGFP-H2B and mCherry-H2B), followed by immunofluorescence (IF) staining with MORC2 and H3K9me3 antibodies (**Fig 4e-f**). Quantification of MORC2 in the histone FRET map shows that MORC2 is associated with significantly lower histone FRET and is localised within the nucleoplasm, indicative of open chromatin regions (**Fig 4g**). In contrast, H3K9me3 IF shows that H3K9me3 is present at the FRET foci, similar to our previous findings in HeLa cells^16^ (**Fig 4h**). This is consistent with our genomic studies, where the majority of MORC2 does not colocalise with H3K9me3.

To ask how MORC2 regulates gene expression, we performed RNAseq analysis on MORC2^WT^ vs MORC2^KO^ (knockout) HEK293T cells (**Extended Data Fig 10e)**, using QL F-tests to compare gene-wise expression levels between MORC2^KO^ and MORC2^WT^ samples. We detected 97 up-regulated genes and 5 down-regulated genes in the MORC2^KO^ compared with the MORC2^WT^ sample (**Extended Data Fig 10f**), indicating that MORC2 is a localised silencer of the genome and does not cause large changes in gene expression. To assess the effect of MORC2^KO^ on chromatin globally, we also perform ATAC-seq on MORC2^KO^ cells. We found that 17 genes were differentially accessible, with 14 genes up-regulated in the MORC2^KO^ sample (**Extended Data Fig 10g)**, including several zinc finger (ZNF) genes that were reported previously^15^. Together, these findings suggest that MORC2 preferentially binds to open chromatin region where H3K9me3 is depleted.

### Phosphorylated MORC2 slows down DNA compaction

Among all MORCs, MORC2 is most similar to MORC1 based on domain organisation. The *C. elegans* MORC1 has been shown to compact DNA^17^, and based on its similarity to MORC2, the same has been proposed for the latter. To test this, we performed a single-molecule DNA compaction fluorescence assay by adding MORC2^WT^, MORC2^1–265^, MORC2^1–603^ or MORC2^PD^ proteins to a double biotinylated λ-DNA immobilised on a coverslip. In the presence of MORC2^WT^ or MORC2^PD^, punctae appeared across the length of λ-DNA confirming that MORC2 compacts DNA (**Fig 5a,b, Supplementary Videos 2, 3**). Applying an in-plane side flow^18^ revealed that these formations were stably condensed clusters (**Fig 5b** and **Supplementary Video 4**), rather than DNA loops. This indicates that MORC2 compacts DNA through a different mechanism than structural maintenance of chromosome (SMC) protein complexes, which achieve compaction by extruding DNA loops^19,20^. Even at lower MORC2 concentrations (0.1 – 5 nM) where DNA compaction transition occurs, MORC2 did not show a DNA loop. The punctae in full-length MORC2 constructs were absent in the CTD lacking MORC2^1–265^ or MORC2^1–603^ constructs, showing that MORC2 CTD is indispensable for its DNA compaction function (**Fig 5c**). Surprisingly, the MORC2^PD^ mutant compacts DNA 3-fold faster than the wild-type (**Fig 5c, Supplementary Video 3**), indicating phosphorylation of PIM motif in MORC2 CTD is crucial for dictating the speed at which DNA compaction would occur.

**Figure 5.**
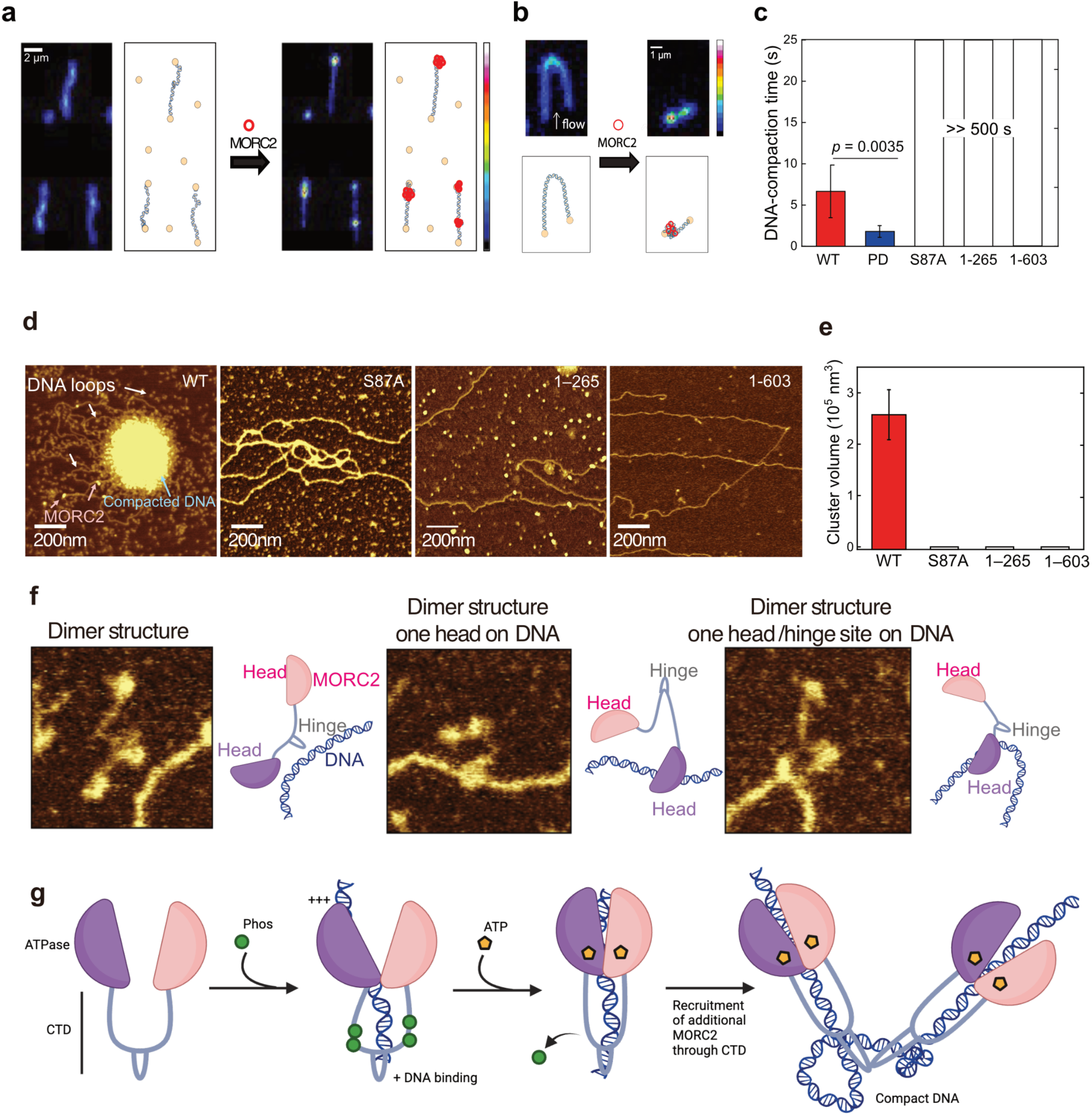
Cluster formation of MORC2 and DNA observed by single-molecule fluorescence imaging and AFM. **a.** Snapshots and schematics of SxO-labeled DNA, before reaction with MORC2 (left) and after (right). Schematics show DNA bound to PEG-slide by biotin-streptavidin. **b.** Snapshot of a side-flow experiment, at the beginning of flow (left) and after compaction (right). **c.** Compaction time of WT, PD, S87A, 1-265 and 1-603 and (mean ± SD, *n* = 11, 6, 20, 20 and 20 images for WT, PD, S87A, 1-265 and 1-603, respectively). **d.** Dry AFM images of WT, S87A, GHKL (1-265) and ATPase (1-603). **e.** Cluster volume with various conditions (n = 2 independent experiments, n = 5 images for each construct). No clusters were observed with MORC2^S87A^, MORC2^1-265^ and MORC2^1-603^. Error bars are based on SEM. **f.** Representative MORC2-mediated DNA binding imaged using dry AFM (n = 2 independent experiments, scale = 0.1 µm x 0.1 µm). Cartoons are schematics of the images of MORC2 dimer at the adjacent of the DNA. **g.** Model of DNA compaction mediated by MORC2 phosphorylation and ATP hydrolysis.

Next, we tested the role of ATP to drive DNA compaction by MORC2. We used an ATPase-dead MORC2^S87A^ construct, which is capable of binding DNA and ATP but could not hydrolyse ATP (**Extended Data Fig 5**). We observed no DNA compaction activity using this construct, hence indicating that ATP hydrolysis is an essential event for MORC2 driven DNA compaction (**Fig 5c, Supplementary Video 5**). To further verify these observations, we used atomic force microscopy (AFM) to image the MORC2-DNA complexes. We found that MORC2^WT^ compacted DNA into a large DNA cluster, whereas the MORC2^S87A^, MORC2^1–265^ and MORC2^1–603^ variants did not (**Fig 5d-e**).The DNA bridging is seen on periphery of the DNA cluster similar to AFM images of yeast cohesin SMC complex in presence of DNA^18^. We next focused on whether DNA binding changes the oligomerisation state of MORC2. We observed a clear colocalization of MORC2 and DNA, with one of the GHKL “head” conformation binding to DNA (**Fig 5f**). Clearly, the clusters were organised by numerous DNA loops, suggesting multiple MORC2 molecules exhibit DNA-trapping behaviour. These results highlight a novel role of MORC2 as an ATP-dependent DNA compaction machine, the efficiency of which is modulated by its phosphorylation.

## Discussion

Dominant variants in MORC2 can cause spinal muscular atrophy and the Charcot-Marie-Tooth (CMT) neuromuscular disease^21^. Further, MORC2 overexpression is associated with several cancers, including breast, gastric and liver cancers^8,10,11^. Notably, MORC2 phosphorylation on serine 739 in its CTD region by PAK1 kinase increases its recruitment to DNA damage sites and causes proliferation of gastric cancer cells^8,9^. In addition to phosphorylation, the MORC2 CTD undergoes other post translational modifications (PTMs) such as acetylation, PARylation and SUMOylation^22,23^. PTMs of CTD are a characteristic feature of all MORC proteins^4,9^. Despite mounting evidence highlighting the importance of MORCs CTD in various cellular processes, there remains a significant gap in our understanding of how the CTD contributes towards the molecular mechanisms underlying chromatin remodelling. Here we undertook a detailed study on MORC2 CTD and explored the functional implications of its phosphorylation state.

Firstly, our genomics and FLIM-FRET analyses showed that MORC2 preferentially binds open chromatin regions, suggesting that DNA, rather than nucleosomes, would be an ideal substrate for *in vitro* studies. MORC2 was previously shown to repress newly formed repetitive element (LINE-1) marked by H3K9me3 to initiate epigenetic silencing^24,25^. Instead, we show that MORC2 is mostly accessible at the promoter region, suggesting that MORC2 plays a broader role poised at promoters to keep transcription in check, consistent with its role as a DNA sliding clamp (**Fig 3e**). Surprisingly, the proteomics analysis of our full-length, insect cell expressed, recombinant MORC2 revealed that the CTD region is heavily phosphorylated, although we do not know which insect cell kinases phosphorylated MORC2. We made a phosphodead construct with mutations of the top 6 phosphorylation sites to test the effect of phosphorylation on MORC2 function. MORC2 constructs containing either phosphorylated or non-phosphorylated CTDs did not induce large structural changes in the ATPase domain upon DNA binding, revealing that the ATPase core is highly stable in the presence of AMP-PNP. The biggest structural changes we observed were rearrangements between the CC1 and CW-CC2 domains based on changes in crosslink clusters in these domains and the increased flexibility of the ATP lid as shown by increased deuterium uptake in this region upon DNA binding. These changes translate into inducing more flexibility in CC1 to drive DNA binding and compaction, which is also evident from the lack of CC1 density in the DNA-bound MORC2 cryoEM structures. The measured DNA binding affinities of full-length MORC2, the ATPase domain, and the phosphodead mutant are similar. However, DNA competition assays indicate that MORC2 holds on to DNA more tightly when its CTD is present. Overall, these results suggest that the CTD enhances MORC2’s ability to stably associate with DNA, which is regulated by its phosphorylation state.

Secondly, we identified multiple DNA binding sites across MORC2. Mechanistically, the low-affinity sites located on the CTD may play a role in genome surveillance in a sequence-independent manner. This suggests that these sites could facilitate a general scanning of the genome similar to what is observed for various transcription factors^26^. We propose that upon receiving a specific signal from the cell to compact a designated region of the genome, MORC2’s high-affinity site traps this region, forming a DNA loop. Following this initial trapping, additional MORC2 molecules are likely recruited via the CTD. In this scenario, the low-affinity sites would transfer the DNA to the high-affinity sites, thereby enabling the further trapping and stabilization of DNA loops (**Fig 5d**). Although we currently lack direct evidence to substantiate this model for MORC2, it is noteworthy that a similar DNA loop-trapping mechanism has been shown for MORC1^17^. However, it is important to note that multiple DNA binding sites have not yet been observed for MORC1. The ability of MORC2 to bind DNA through its multiple sites, combined with the flexibility of its CTD relative to the ATPase domain, explains why it is hard to determine DNA-bound MORC2 structures. This likely explains the absence of full-length and/or DNA-bound structures for MORCs in general, indicating similar structural challenges for other MORC proteins^2,3^.

Finally, we provided direct visual evidence for DNA binding and compaction by MORC2. Both the full-length wild type (WT) and phosphodead mutant of MORC2 efficiently compacts DNA, whereas the ATPase domain lacking the CC1 (1-265) and CC3-CC4 domain (1-603) do not, demonstrating the indispensable nature of the CTD in MORC2’s function. The phosphodead mutant compacts DNA three times faster than the WT, which correlates with its two-fold higher ATP hydrolysis rate. We tested the full-length MORC2 with the patient variant S87L, which binds ATP but exhibits significantly lower ATPase activity (**Extended Data Figure 5e**), consistent with previous findings^7^. Based on this patient variant, we designed a S87A mutant that binds DNA and ATP but cannot hydrolyse ATP. The S87A mutant binds to AMP-PNP but does not compact DNA (**Extended Data Fig 5b**, **Fig 5c**), highlighting the critical role of ATP hydrolysis in this process. This contrasts with *Caenorhabditis. elegans* MORC1, which only requires ATP binding for DNA compaction^17^. We provided direct evidence, using side-flow single-molecule experiments, that MORC2 does not extrude a DNA loop. Instead of observing gradual loop growth, we observed DNA cluster formation (**Fig 5b**), similar to DNA compaction findings for MORC1^17^. Given the structural similarities between MORC2 and MORC1, and the limited gene expression changes observed upon MORC2 knockout, it is likely that MORC2 compacts DNA by loop trapping on specific DNA regions, such as the PARP1-induced DNA breaks or HUSH-associated H3K9me3 marks^15,22^. This contrasts with the global chromatin compaction by canonical SMC protein complexes, which extrudes supercoiled DNA to alter the 3D shape of the genome^27^. MORC1 can compact chromatinised DNA, but at a slower rate than naked DNA^17^, so we are intrigued to see whether MORC2 would do the same. Our findings establish ATP hydrolysis-driven DNA compaction as a major function of MORC2. Future research should explore how patient mutations in MORC2, which alter ATP hydrolysis activity, impact its DNA compaction ability.

While this work provides mechanistic insights into how CTD and its phosphorylation modulates MORC2 function, several questions remain. For instance, why does the majority of MORC2 occupy promoter regions in cells without external stimuli? How is MORC2 recruited to H3K9me3-marked heterochromatin regions? And how does MORC2 switches its role between DNA damage response and epigenetic silencing? Additionally, the effect of other PTMs on MORC2 activity at the molecular level needs further exploration. Addressing these questions will require a concerted effort integrating multiple technologies.

In conclusion, our work shows that the phosphorylation of MORC2 CTD fine tunes its DNA compaction activity. Important findings include (1) our demonstration that full-length MORC2 is an active dimer that can be stabilised with ATP binding, reopening the investigation for MORC2 substrates for chromatin remodelling; (2) removal of phosphorylation at conserved coiled coil domains increases the rate of ATP hydrolysis, which will recycle MORC2 to proximal and adjacent DNA substrates; and (3) regulation of DNA compaction regulation by ATP hydrolysis. Our mechanistic insights will help efforts to develop therapeutic strategies targeting MORC2 to target to treat various neurological disorders and cancers.

## Methods

### Protein expression and purification

Full-length human MORC2 (UniProt id: Q9Y6X9) and PAK1 (UniProt id: Q13153) cDNAs were synthesized by Epoch Life Sciences. For protein expression, full-length MORC2 and its variants (residues 1–265, 1–496, 496–603, 496–1032, 604–1032, 544-587 (CC2), 738-776 (CC3) 963-1020 (CC4), S87A, PD(S725A, S730A, S739A, S743A, S777A, S779A), S87L) and PAK1 were sub-cloned into a pACEBac1 vector with a C-terminal 3C protease-cleavage site followed by twin strepII-tag. MORC2 ATPase (residues 1-603) was cloned into pFastbac construct with 6xHis at the N-terminus and purified as previously described^13^. Each construct was transformed into EMBacY cells to generate a bacmid. Bacmid DNA was transfected into Sf9 cells using FuGENE transfection reagent (Promega) and virus was passaged twice in the same cell line before large-scale infection. Sf9 cells were a kind gift from the Glukhova lab at WEHI (negative for mycoplasma, identity not independently authenticated by us). For large-scale expression, Sf9 cells at a density of 2–2.5 × 10^6^ cells per mL were infected with 1% (v/v) second passage (P2) virus and incubated at 27 °C for 50–60 h, 220 rpm.

The cells were harvested by centrifugation, resuspended in lysis buffer containing 50 mM HEPES, pH 8.0, 300 mM NaCl, 1 mM TCEP, 5% glycerol, 10 U per ml benzonase solution (Sigma), 1× cOmplete EDTA-free protease inhibitors (Roche) and 5 mM benzamidine hydrochloride. Cells were lysed by sonication. Clarified cell lysate was incubated with BioLock (IBA Lifesciences) and Strep-Tactin resin (IBA Lifesciences) for 1 h followed by wash with lysis buffer. All MORC2 constructs (except 1-495 and 495-603) and PAK1 proteins were eluted in elution buffer (100 mM HEPES pH 8.0, 300 mM NaCl, 1 mM TCEP, 5% glycerol and 8 mM desthiobiotin). Further purification was performed by 1 mL HiTrap Q Sepharose Fast Flow column (Cytiva) using a linear gradient of NaCl from concentration of 150 mM to 1 M in 50 mM HEPES pH 8.0, 1 mM TCEP, over 22 column volumes. For MORC2 1-495 and 495-603, this was followed by Heparin affinity chromatography (GE Healthcare) in the same buffer using a linear gradient of NaCl from concentration of 150 mM to 1 M over 20 column volumes. The final buffer for purified MORC2 constructs was 50 mM HEPES pH 8.0, ∼500 mM NaCl, 1 mM TCEP. The fractions containing MORC2 were pooled and concentrated using 10 kDa or 30 kDa MWCO Vivaspin Concentrator (Biostrategy), and further purified on size exclusion chromatography using the Superose 6 Increase 10/300 GL column (Cytiva).

### Size exclusion chromatography multi-angle light scattering (SEC-MALS)

Purified proteins were run on an XBridge Protein BEH SEC Column (Waters) coupled with DAWN HELEOS II light scattering detector and Optilab T-rEX refractive index detector (Wyatt Technology). The system was equilibrated in 50 mM HEPES pH 8.0, 300 mM NaCl, 1 mM TCEP and calibrated using bovine serum albumin (2 mg ml^−1^) before analysis of experimental samples. For each experiment, 20 µl of purified protein (1 mg ml^−1^) was injected onto the column and eluted at a flow rate of 0.5 ml min^−1^. Experimental data were collected and processed using ASTRA (Wyatt Technology, v.7.3.19).

### Electrophoretic mobility shift assay (EMSA)

Different DNA substrates were formed by annealing oligonucleotides (synthesised by IDT, described in **Supplementary Table 5**) at 95 °C for 5 min and slowly cooling down to room temperature over ∼2 h. The X0m1 oligonucleotide was 6FAM-, IRDye680- or IRDye800-labelled at 5’ end. The Xom1 or P1 oligonucleotides were annealed with their corresponding X0m1.com or P7 oligonucleotides. For all DNA binding experiments, 25 nM DNA was incubated in 10 µL reaction buffer (20 mM Tris pH 7.4, 75 mM NaCl, 5% glycerol, 0.005% NP-40, 0.5 mM MgCl_2_) containing increasing concentrations of MORC2 (0, 25, 50, 100 nM) at room temperature for 30 min. For competitive EMSA experiments, 25 nM IRDye680-dsDNA (60bp) or cirDNA (105bp) was incubated with 100 nM of MORC2 proteins at room temperature for 30 min, followed by addition of competitive IRDye800-dsDNA (60bp) with increasing concentrations (0, 25, 100, 400, 2000 nM) at room temperature for 10 minutes. The reaction was resolved by electrophoresis on a 6% non-denaturing polyacrylamide gel in TBE (100 mM Tris, 90 mM boric acid, 1 mM EDTA) buffer and visualised by ChemiDoc (BioRad) using SYBR Gold stain or imaging in the 6FAM, IRDye680 and IRDye800 channels.

### *In vitro* kinase and phosphatase assay

Ten μg of purified MORC2 and 0.1 μg of purified PAK1 were incubated in 60 μL of reaction buffer (20 mM Tris-HCl pH 7.4, 10 mM MgAc, 0.5 mM DTT, 0.05% Tween-20, 100 mM KCl and 0.2 mM ATP) for 30 min at 30 °C. For phosphatase experiments, 600 units of lambda phosphatase (NEB) was added to reactions together with 1 mM MnCl2 and incubated for 30 min at 30 °C. All kinase reactions were quenched in 1X Laemmli sample buffer (Bio-rad) and run on 12.5% SuperSep Phos-tag gels (FujiFilm) containing 25 mM Tris, 250 mM glycine and 0.1% SDS running buffer. Gels were stained with InstantBlue Coomassie Protein Stain (Abcam). Gel bands containing phosphorylated MORC2 were excised for mass spectroscopy analysis.

### Mass spectrometry-based proteomics

Protein gel bands were carefully excised and were destained in a 50% acetonitrile (ACN) solution for 30 minutes at 37°C. Subsequently, the gel band was dehydrated using 100% ACN at room temperature for 15 minutes. The ACN was aspirated, and the gel piece was further dried using a vacuum centrifuge (CentriVap, Labconco).

Proteins were reduced using 1 mM dithiothreitol (DTT) in 50 mM ammonium bicarbonate for 30 minutes at RT. Any excess DTT was carefully aspirated before adding 55 mM iodoacetamide to perform alkylation. Alkylation was performed at RT for 60 minutes. Following reduction and alkylation, the gel piece was washed again using 50% ACN twice, followed by a single wash with 100% ACN and drying the gel piece again.

Proteins within the gel piece were then subjected to overnight digestion using 500 ng trypsin in 50 mM ammonium bicarbonate, carried out at a temperature of 37 degrees. The following day, peptides were extracted using a solution of 60% ACN and 0.1% formic acid (FA). The collected peptides were subsequently lyophilised to dryness using a CentriVap (Labconco). The final step involved reconstituting the peptides in a 10 µl solution containing 0.1% FA and 2% ACN, preparing them for analysis via mass spectrometry.

The digested peptides were reconstituted in a solution containing 2% ACN and 0.1% formic acid (FA), and subsequently analysed using an Orbitrap Eclipse Tribrid mass spectrometer coupled with a Neo Vanquish LC. The samples were loaded onto a C18 fused silica column (inner diameter 75 µm, outer diameter 360 µm, and length 15 cm, with 1.6 µm C18 beads) packed into an emitter tip (IonOpticks) using pressure-controlled loading, with a maximum pressure of 1,500 bar. The system was interfaced to an Orbitrap Eclipse™ Tribrid mass spectrometer via Easy nLC source, and the peptides were electrosprayed directly into the mass spectrometer.

For liquid chromatography (LC) separation, a linear gradient of solvent-B (99% acetonitrile) was applied at a flow rate of 400 nl/min. The gradient started at 2% solvent-B and increased to 30% over 5 minutes, followed by a gradient from 5% to 17% solvent-B for 15 minutes and from 17% to 34% solvent-B for the next 28 minutes. A column washing step at 85% solvent-B for 9 minutes concluded the 54-minute. Data was acquired in in a data-dependent acquisition (DDA) mode.

During MS1 acquisition, spectra were obtained in the Orbitrap with a resolution of 120,000 (R), standard normalized automatic gain control (AGC) target, Auto MaxIT, RF Lens set at 30%, and a scan range of 380–1400. Dynamic exclusion was implemented for 30 seconds, excluding all charge states for a given precursor. Data-dependent MS2 spectra were collected in the Orbitrap for precursors with charge states 3-8, utilizing a resolution of 50,000 (R), assisted HCD collision energy mode, normalized HCD collision energies set at 25% and 30%, normal scan range mode, normalized AGC target of 200%, and MaxIT of 150 ms.

The raw mass spectrometry (MS) data files underwent processing using MaxQuant version 2.0.1.0, incorporating the Andromeda search engine [23]. The analysis was conducted against the Human proteome fasta database obtained from UniProt in April 2022. Variable modifications included protein N-terminal acetylation, methionine oxidation, and phosphorylation on serine, threonine, and tyrosine (STY), while cysteine carbamidomethylation was selected as a fixed modification. Trypsin served as the specified enzyme for digestion, with an allowance for up to two missed cleavages. To ensure high confidence in the results, the false discovery rate (FDR) was set at 1% for both proteins and peptides. A peptide tolerance of 4.5 ppm was used during main search.

### Cryo-electron microscopy (cryoEM)

Purified proteins were vitrified by applying 3-4 µL purified protein (2 µM), either in presence of three-fold molar excess of 60 bp DNA or in its absence, to UltrAuFoil R1.2/1.3 grids (Quantifoil). The grids were glow discharged in the presence of amylamine in PELCO easiGlow. The grids were prepared using a Vitrobot Mark IV (ThermoFisher) at 4°C and 100% humidity with blotting force of −3 N for 3 s. The MORC2^S87A^ and MORC2^PD^ grids were imaged on Titan Krios microscope operated at 300 keV using a Gatan K3 detector. The MORC2^1–603^ grids were collected on Talos Arctica operating at 200 keV using a Gatan K2 detector.

### CryoEM image processing and model building

Image processing was performed in cryoSPARCv3 and v4^28^. For all datasets, the movies were aligned and averaged using patch motion correction, and contrast transfer function parameters were estimated using Patch CTF estimation. Particles were picked using crYOLO^29^, and the particle coordinates were transferred to cryoSPARC for 2D class average. After 2D classification of the auto-picked particles, selected classes were used to make an ab initio model. Heterogenous refinement with a ‘junk’ model was used to clean up particles for higher resolution template. The particles were further refined using homogenous refinement protocol with imposed C2 symmetry. Final set of particles reaching 1.9 – 3.2 Å were used to perform non-uniform refinement with imposed C2 symmetry. The 3D maps were post-processed to automatically estimate and apply the B-factor and to determine the resolution by Fourier shell correlation between two independent half datasets using 0.143 criterion. Local resolution was estimated using CryoSparc “Local Resolution Estimation” Tool (version 4.5.1). Directional FSC and sphericity values for the maps were generated using the 3DFSC server (https://3dfsc.salk.edu/)^30^.

The crystal structure of MORC2 ATPase domain (PDB: 5OF9) was used for modelling into MORC2^PD^, MORC2^1–603^, MORC2^PD-DNA^ and MORC2^1–603-DNA^ and the crystal structure of MORC2 S87L variant (PDB: 5OFB) was used for MORC2^S87A^. Model placement and initial refinement was performed in Coot^31^ and further refined using Phenix refinement^32^. ChimeraX (v.1.7) was used to color the surface of the cryoEM maps based on local resolution parameters obtained from Cryosparc. All images were rendered in ChimeraX (v.1.7).

### Generation of MORC2 KO HEK293 cells

Two guide RNAs (gRNAs) were designed using Synthego for human MORC2 (https://design.synthego.com/%23/) to generate two biological bulk replicates. The gRNA sequences are UUCAGGGGCUCAAUGCGCAU and UCAGGGGCUCAAUGCGCAUU. These gRNA were ordered from IDT. The SF cell line 4D-Nucleofector X Kit S (V4XC-2032) from Lonza was used to prepare a mix of 300 pmol of gRNA and 100 pmol of Cas9 protein. About 1 × 10^6^ HEK293 cells were electroporated using Lonza Amaxa TM machine following the manufacturer’s instruction. The transfected cells were cultured until they become confluent and split two more times before freezing down. The KO efficiency was measured by performing western blot analysis using anti-MORC2 antibody (PA551172, **Extended Data Fig 11a**) on the cell lysate (∼10,000 cells) from both the gRNAs. The gRNA1 and gRNA2 KO cells were used for our downstream assays. RNAseq, ATAC-seq and ChIP-seq were each performed on 2 independent WT replicates, and 2 independent KO cell lines generated using different MORC2-targeting sgRNA sequences.

### Fluorescence polarization ATPase assay

Fluorescence polarization ATPase assays were performed as outlined in^33^. A 10 mL reaction was set up in triplicates in 384-well low flange, black, flat-bottom plates (Corning) containing 7 mL reaction buffer (50 mM HEPES pH 7.5, 4 mM MgCl_2_, 2 mM EGTA), 1 mL recombinant protein at concentrations ranging from 0.1-0.6 mM or SEC buffer control, 1 mL nuclease-free water and 1.25-10 mM ATP substrate. Reactions were incubated at 25 °C for 1 hour in the dark. Reactions were stopped by the addition of 10 mL detection mix (1X Detection buffer, 4 nM ADP Alexa Fluor 633 Tracer, 128 mg/mL ADP^2^ antibody) and incubated for another hour in the dark. Fluorescence polarization readings (mP) were measured using an Envision plate reader (PerkinElmer Life Sciences) fitted with excitation filter 620/40 nm, emission filters 688/45 nm (s and p channels) and D658/fp688 dual mirror. Readings from a free tracer (no antibody) control were set as 20 mP as the normalization baseline of the assay for all reactions. The amount of ADP produced by each reaction was estimated by a 12-point standard curve, as outlined in the manufacturer’s protocol. Data were plotted and analyzed in GraphPad Prism.

### Surface plasmon resonance (SPR)

SPR binding studies of MORC2 variants to DNA were performed using a Biacore S200 Instrument (Cytiva). Biotinylated 60 bp dsDNA was prepared by assembling oligonucleotides 1 (/5Biosg/ACG CTG CCG AAT TCT ACC AGT GCC TTG CTA GGA CAT CTT TGC CCA CCT GCA GGT TCA CCC) and oligonucleotides 2 (GGG TGA ACC TGC AGG TGG GCA AAG ATG TCC TAG CAA GGC ACT GGT AGA ATT CGG CAG CGT). Biotinylated 60 bp dsDNA was diluted to 5 µg/mL in SPR running buffer (10 mM HEPES pH 7.4, 300 mM NaCl, 3 mM EDTA and 0.05% (v/v) NP-20) to a final immobilisation level of 200-220 response units (RU) on the Streptavidin chip (Cytiva). An additional injection step with 1 min of Biolock (IBA) was used to block StrepII-tagged MORC2 from binding to the Streptavidin chip. A blank activation/deactivation was used for the reference surface. DNA binding studies were performed at 20 °C in SPR running buffer. MORC2^FL^, MORC2^S87A^, MORC2^1–495^, MORC2^1–603^, and MORC2^496–1032^ were diluted to 500 nM stock in SPR running buffer and prepared as a 8-point concentration series (2-fold serial dilution, 7-500 nM). Samples were injected in a multi-cycle run (flow rate 30 µL/min, contact time of 60 s, dissociation 120 s) with regeneration with 0.5 M EDTA buffer. Sensorgrams were double referenced, and steady-state binding data fitted using a 1:1 binding model using Biacore S200 Evaluation Software (Cytiva, v1.1). Representative sensorgrams and fitted dissociation constant (*K*_D_) values, depicted as mean ± SEM (n>3 independent experiments) are shown in Fig 3a.

### Quantitative Crosslinking Mass Spectrometry

Sulfo-SDA (ThermoFisher) was dissolved freshly in assay buffer (50 mM HEPES pH 8.0, 300 mM NaCl, 1 mM TCEP) to a final concentration of 0.5 mg/mL. Crosslinking reactions were performed in triplicate by incubating 0.5 mg/mL MORC2 in the presence of absence of AMP-PNP and DNA, with 0.5 mg/mL sulfo-SDA in a final volume of 10 µL at room temperature for 30 min. To activate sulfo-SDA, the samples were irradiated with a 1000 kV UV lamp at 356 nm on ice for 1 min. Crosslinked samples were analysed on a 3-8% NuPAGE Tris-acetate gel (Invitrogen), and gel bands corresponding to the crosslinked MORC2 were excised and digested as previously described above^34^.

Reconstituted peptides were analysed on Orbitrap Eclipse Tribrid mass spectrometer that is interfaced with Neo Vanquish liquid chromatography system. Samples were loaded on to a C18 fused silica column (inner diameter 75 µm, OD 360 µm × 15 cm length, 1.6 µm C18 beads) packed into an emitter tip (IonOpticks) using pressure-controlled loading with a maximum pressure of 1,500 bar, that is interfaced to an Orbitrap Eclipse™ Tribrid (Thermo Scientific) mass spectrometer using Easy nLC source and electro sprayed directly into the mass spectrometer.

We then employed a linear gradient of 3% to 30% of solvent-B at 400 nl/min flow rate (solvent-B: 99% (by vol) acetonitrile) for 100 min and 30% to 40% solvent-B for 20 min and 35% to 99% solvent-B for 5 min which was maintained at 90% B for 10 min and washed the column at 3% solvent-B for another 10 min comprising a total of 145 min run with a 120min gradient in a data dependent (DDA) mode. MS1 spectra were acquired in the Orbitrap (R = 120k; normalised AGC target = standard; MaxIT = Auto; RF Lens = 30%; scan range = 380–1400; profile data). Dynamic exclusion was employed for 30 s excluding all charge states for a given precursor. Data dependent MS2 spectra were collected in the Orbitrap for precursors with charge states 3-8 (R = 50k; HCD collision energy mode = assisted; normalized HCD collision energies = 25%, 30%; scan range mode = normal; normalised AGC target = 200%; MaxIT = 150 ms).

Quantitative crosslinking analysis was performed following the previously published method^34^. Raw data files were converted to MGF files using MS convert^35^. MGF files were searched against a fasta file containing the MORC2 sequence using XiSearch software^36^ (version 1.7.6.7) with the following settings: with the following settings: crosslinker = multiple, SDA and noncovalent; fixed modifications = Carbamidomethylation (C); variable modifications = oxidation (M), sda-loop (KSTY) DELTAMASS:82.04186484, sda-hydro (KSTY) DELTAMASS:100.052430; MS1 tolerance = 6.0ppm, MS2 tolerance = 20.0ppm; losses = H_2_O,NH_3_, CH_3_SOH, CleavableCrossLinkerPeptide:MASS:82.04186484). False Discovery Rate (FDR) was performed with the in-built xiFDR set to 5%^37^. Identified crosslinks were converted to linear peptide sequences using the skyline convert tool and quantities were calculated with Skyline^34^. Data were visualised using the XiView software ^38^.

Data cleaning, pre-processing and statistical analysis were conducted using R (version 4.2.1). Decoy precursors and cross-linked peptides with Isotope.Dot.Product < 0.8 were filtered out. Additionally, cross-linked peptides that quantified in at least 67% of replicates within at least one condition were considered for further analysis. Total of 910 cross-linked peptides were included in the analysis. The normalised area of cross-linked peptides were log-transformed and missing values were imputed using the Barycenter approach (v2-MNAR) as implemented in the msImpute package (v.1.8.0)^39^. Multivariate analysis, principal component analysis (PCA), was employed to identify any potential outliers. Differential analysis was carried out using the limma package (v. 3.54.2)^40^. A cross-linked peptide was deemed significantly differentially expressed if the FDR was ≤ 5% following Benjamini–Hochberg (BH) correction.

### Hydrogen deuterium exchange mass spectrometry

MORC2 ATPase domain (1-603) and CTD (496-1032) were assayed with 60 bp dsDNA (1:3 molar ratio of protein to DNA) or in absence of DNA. All samples had 2.5 mM AMP-PNP. HDX labelling of protein was performed at 20°C for periods of 6, 60, 600, 6000 s using a PAL Dual Head HDX Automation manager (Trajan/LEAP) controlled by the ChronosHDX software (Trajan). A 4 µL of the sample (containing ∼15 µM of protein) was transferred to 55 µL of non-deuterated (50 mM potassium phosphate buffer pH 7.4, 150 mM NaCl, in H_2_O) or deuterated (50 mM potassium phosphate buffer pD 7.0, 150 mM NaCl, in D_2_O) buffer and incubated for the respective time. Quenching was performed by adding 50 µL of the deuterated protein to 55 µL of quench buffer (50 mM potassium phosphate buffer, pH 2.3 containing 4 M guanidine hydrochloride) at 1°C. For online pepsin digestion, 95 µL of the quenched sample was passed over an immobilized 2.1 × 30 mm Enzymate BEH pepsin column (Waters) equilibrated in 0.1% (v/v) formic acid in water at 100 µL/min. Proteolyzed peptides were captured and desalted by a C18 trap column (VanGuard BEH; 1.7 μm; 2.1 × 5 mm (Waters)) and eluted with acetonitrile and 0.1% (v/v) formic acid gradient (5% to 40% in 8 min, 40% to 95% in 0.5 min, 95% 1.5 min) at a flow rate of 40 μL/min and separation on an ACQUITY UPLC BEH C18 analytical column (1.7 μm, 1 × 100 mm (Waters)) delivered by ACQUITY UPLC I-Class Binary Solvent Manager (Waters).

For mass spectrometry, a SYNAPT G2-Si mass spectrometer (Waters) was used. Instrument settings were: 3.0 KV capillary and 40 V sampling cone with source and desolvation temperature of 100 and 40°C respectively. The desolvation and cone gas flow was at 80 L/hr and 100 L/hr, respectively. High energy ramp trap collision energy was from 20 to 40 V. All mass spectra were acquired using a 0.4 s scan time with continuous lock mass (Leu-Enk, 556.2771m/z) for mass accuracy correction. Data were acquired in MS^E^ mode.

Raw data obtained from non-deuterated samples were processed using Protein Lynx Global Server (PLGS) v3.0 (Waters). This processing used a database that contained sequence information about *Sus scrofa* pepsin A and each MORC2 construct. PLGS processing parameters were 135 and 30 counts for low and elevated energy threshold, respectively. Primary digest reagent was set to non-specific and with oxidation methionine and phosphorylation of serine, threonine and tyrosine as variable modification. FDR was set to 1%. In the case of labelled data, the analysis was performed using DynamX 3.0, with 5000 minimum intensity criteria, 0.3 products per amino acid, 1 consecutive product, and error of 10 ppm. The file threshold was 3 out of 4. Data were manually curated. Heatmaps were made using in-house Python scripts (D’Arcy laboratory). Structural mappings were done using scripts from DynamX in PyMOL version 2.5.

### Fluorescence Microscopy

#### Microscopy data acquisition

All microscopy measurements were performed on an Olympus FV3000 laser scanning microscope coupled to an ISS A320 Fast FLIM box for fluorescence fluctuation data acquisition. A 60X water immersion objective 1.2 NA was used for all experiments and the cells were imaged at 37 degrees in 5% CO2. For single channel Number and Brightness (NB) measurement the eGFP tagged plasmids were excited by a solid-state laser diode operating at 488 nm and the resulting fluorescence signal directed through a 405/488/561 dichroic mirror to a photomultiplier detector (H7422P-40 of Hamamatsu) fitted with an eGFP 500/25 nm bandwidth filter. NB fluorescence fluctuation spectroscopy (FFS) measurement of the eGFP tagged MORC2 construct involved selecting a HEK293T MORC2 KO cell exhibiting low eGFP expression level which enabled observation of fluctuations in eGFP fluorescence intensity and then selecting a 10.6 μm region of interest (ROI) within that MORC2 KO cell’s nucleus, which for a 256 x 256 pixel frame resulted in a pixel size of 41 nm (an oversampling of the point spread function of our diffraction limited acquisition). A frame scan acquisition (*N* = 100 frames) was then acquired with the pixel dwell time set to 12.5 µs, which resulted in a line time of 4.313 ms and a frame time of 1.108 s. HEK293T nucleus co-expressing the histone FRET pair H2B-eGFP and H2B-mCH that has been fixed with IF against MORC2 or H3K9me3. The FLIM imaging involved sequential imaging of a two-phase light path in the Olympus FluoView software. The first phase was set up to image H2B-eGFP and H2B-mCh via use of solid-state laser diodes operating at 488 and 561 nm, respectively, with the resulting signal being directed through a 405/488/561/6033 dichroic mirror to two internal GaAsP photomultiplier detectors set to collect 500–540 nm and 600–700 nm. The second phase was set up to image AF647 via use of solid-state laser diodes operating at 633 nm, with the resulting signal being directed through a 405/488/561/633 dichroic mirror to the internal GaAsP photomultiplier detector set to collect 600–700 nm. Then in each HEK293T nucleus selected, a FLIM map of H2B-eGFP was imaged within the same field of view (256×256-pixel frame size, 20 µs/pixel, 90 nm/pixel, 20 frame integration) using the ISS VistaVision software. This involved excitation of H2B-eGFP with the external pulsed 488 nm laser (80 MHz) and the resulting signal being directed through a 405/488/561/633 dichroic mirror to an external photomultiplier detector (H7422P-40 of Hamamatsu) that was fitted with a 520/50 nm bandwidth filter. The donor signal in each pixel was then subsequently processed by the ISS A320 FastFLIM box data acquisition card to report the fluorescence lifetime of H2B-eGFP. All FLIM data were pre-calibrated against fluorescein at pH 9 which has a single exponential lifetime of 4.04 ns.

#### Number and brightness (NB) analysis

The brightness of a fluorescently tagged protein is a readout of that protein’s oligomeric state that can be extracted by a moment-based Number and brightness (NB) analysis of an FFS frame scan acquisition. In brief, within each pixel of a frame scan we have an intensity fluctuation that has an average intensity (first moment) and a variance (second moment). As defined in Equation 1, the ratio of these two properties describes the apparent brightness (B) of the molecules that give rise to the intensity fluctuation.

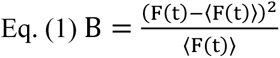

The true molecular brightness (*ɛ*) of the molecules is related to the measured apparent brightness (*B*) by, where 1 is the brightness contribution of our photon counting detector. Calibration of the apparent brightness of monomeric eGFP (B_monomer_ = 1.15 ± 0.05) enabled extrapolation of the expected apparent brightness of eGFP-HP1α dimers (B_dimer_ = 1.30 ± 0.1) and oligomers (B_oligomer_ = 1.60 ± 0.2), which in turn enabled definition of brightness cursors to extract and spatially map the fraction of pixels within a given frame scan acquisition that contain these different species. The fraction of eGFP-HP1α dimer and eGFP-HP1α oligomer (i.e., number of pixels assigned B_dimer_ or B_oligomer_) were used as parameters to quantify the degree of HP1α self-association detected across multiple cells. Artefacts due to cell movement or photobleaching were subtracted from acquired intensity fluctuations via use of a moving average algorithm. All brightness calculations were carried out in SimFCS from the Laboratory for Fluorescence Dynamics (www.lfd.uci.edu).

#### FLIM-FRET Analysis

The fluorescence decay recorded in each pixel of a FLIM image was quantified by the phasor approach to lifetime analysis. Each pixel of the FLIM image gives rise to a single point (phasor) in the phasor plot, and, when used in reciprocal mode, enables each point of the phasor pot to be mapped to each pixel of the FLIM image. Since phasors follow simple vector algebra, it is possible to determine the fractional contribution of two or more independent molecular species coexisting in the same pixel (e.g., autofluorescence and a fluorescent protein). In the case of two independent species, all possible weightings give a phasor distribution along a linear trajectory than joins the phasors of the individual species in pure form. In the case of a FRET experiment where the lifetime of the donor molecule is changed upon interaction with an acceptor molecule, the realization of all possible phasors quenched with different efficiencies describe a curved trajectory in the phasor plot. The FRET trajectory follows the classical definition of FRET efficiency. As described in previous papers^16,41^, the contribution of the background (cellular auto-fluorescence) and of the donor without acceptor were evaluated by using the rule of the linear combination, with the background phasor and donor unquenched determined independently. By linking two phasor cursors between these two terminal phasor locations, we can pseudo-colour each pixel of a FLIM map according to the exact FRET efficiency detected at that location. Furthermore, from linking two additional cursors between the donor phasor and background phasor, we can also quantify the contribution of cellular autofluorescence in each pixel. To increase phasor accuracy, a 3 x 3 spatial median filter was applied twice to the FLIM maps presented in before FRET analysis. All FLIM-FRET quantification was performed in the SimFCS software developed at the LFD (www.lfd.uci.edu).

#### Immuno Fluorescence (IF) mask analysis of histone FRET

To quantify the chromatin nanostructure associated with different MORC2 or H3K99me3, we applied an intensity threshold mask based on IF intensity images to FLIM maps pseudo-coloured according to histone FRET (compact chromatin) versus no FRET (open chromatin). This involved: (1) smoothing each HEK293T nucleus’ IF image with a 3×3 spatial median filter, (2) transforming this smoothed image into a binary mask based on an intensity threshold that retains the top ∼5% intensity pixels, (3) applying the IF-guided mask to its associated FLIM map pseudo-coloured according to histone FRET, and (4) quantification of the fraction of compact chromatin within high intensity MORC2/H3K9me3 region versus outside the IF-guided mask (nucleoplasm) in total.

### ChIP-seq

ChIP-seq was performed as per^42^ with amendments. Briefly, 5-10 × 10^6^ cells per IP were cross-linked by adding 1/10 volume of fresh 11% formaldehyde solution (50mM HEPES-KOH pH 7.4, 100 mM NaCl, 1 mM EDTA, 1 mM EGTA, 11% formaldehyde) to a 1 × 10^6^ cell suspension for 20 minutes at room temperature with mixing. 1/12.5 volumes of 2.5 M glycine was added to quench formaldehyde and cells were washed twice in ice-cold PBS. Cross-linked cells were lysed in Lysis Buffer 1 (50 mM HEPES-KOH pH 7.5, 140 mM NaCl, 1 mM EDTA, 10% (v/v) glycerol, 0.5% (v/v) IGEPAL CA-630, 0.25% (v/v) Triton X-100, 1x Roche cOmplete™, EDTA-free Protease Inhibitor Cocktail) for 10 minutes at 4 °C with rocking, pelleted at 1350 x *g* for 5 minutes at 4°C, then Lysis Buffer 2 (10 mM Tris-HCl pH 8.0, 200 mM NaCl, 1 mM EDTA, 0.5 mM EGTA, 1x Roche cOmplete™, EDTA-free Protease Inhibitor Cocktail) for 10 minutes at room temperature with rocking, pelleted again at 1350 x *g* for 5 minutes at 4 °C, then resuspended in Lysis Buffer 3 (10 mM Tris-HCl pH 8.0, 100 mM NaCl, 1 mM EDTA, 0.5 mM EGTA, 0.1% (w/v) Na-Deoxycholate, 0.5% (v/v) N-lauroylsarcosine, 1x Roche cOmplete™, EDTA-free Protease Inhibitor Cocktail) and transferred to Covaris miniTUBEs. Cell lysates were sonicated using a Covaris E220 focused-ultrasonicator with the following settings to shear DNA to a size of 300-500 bp suitable for high-throughput sequencing: Fill Level 10, Duty Cycle 10, PIP 175, Cycles/Burst 200 for 80 seconds. 1/10 volume of 10% (v/v) Triton X-100 was then added to the sonicated lysate and cellular debris was pelleted at 20,000 x *g* for 10 minutes at 4 °C. Immunoprecipitation was then performed overnight at 4 °C with rotation using ProteinG Dynabeads (50 μL per IP) pre-coupled with antibody (5 μL per IP) in 0.5% BSA/PBS for >5 h. Antibodies used were anti-MORC2 Rabbit pAb (Invitrogen #PA5-51172), H3K9me3 Rabbit pAb (Abcam #8898), Rabbit IgG pAb (Abcam #ab46540). Following immunoprecipitation, beads were collected using a magnet and washed 5 times with Wash Buffer (50 mM HEPES-KOH pKa 7.55, 500 mM LiCl, 1 mM EDTA, 1% (v/v) IGEPAL CA-630, 0.7% (w/v) Na-Deoxycholate, then 1 time with Tris-EDTA-NaCl buffer (10 mM Tris-HCl pH 8.0, 1 mM EDTA, 50 mM NaCl). DNA was then eluted from beads by incubation in Elution Buffer (50 mM Tris-HCl pH 8.0, 10 mM EDTA, 1% (w/v) SDS) at 65 °C for 30 minutes with shaking. Bead-free supernatant was reverse crosslinked by incubation at 65 °C for 6-15 h, diluted with equal volume of TE Buffer (10 mM Tris-HCl pH 8.0, 1 mM EDTA), then digested sequentially with RNaseA (0.2 mg/mL) at 37 °C for 2h and Proteinase K (0.2 μg/mL) at 55 °C for 2 h. Final immunoprecipitated DNA was then purified using Zymo ChIP DNA Clean & Concentrator kit and prepared for sequencing using Illumina Truseq ½ reaction library preparation kit. Final libraries were quantified using an Agilent Tapestation using a D1000 screentape and sequenced using an Illumina NextSeq2000 to generate libraries with 24 – 119 million paired-end 65 bp reads.

### ATAC-seq

ATAC-seq was performed using the Omni-ATAC method as described previously^43^. Briefly, 50,000 cells were pelleted at 500 x *g* for 5 minutes at 4 °C then lysed on ice for 3 minutes in 50 μL ATAC-RSB Buffer (10mM Tris-HCl pH 7.4, 10mM NaCl, 3mM MgCl_2_) also containing 0.1% (v/v) IGEPAL CA-630, 0.1% (v/v) Tween-20, 0.01% (w/v) Digitonin. After lysis, 1mL of ATAC-RSB containing only 0.1% (v/v) Tween-20 but no IGEPAL CA-630 or Digitonin was added, and nuclei were pelleted at 500 x *g* for 10 minutes at 4 °C. All supernatant was aspirated and discarded, and each pellet of nuclei was resuspended in 50 μL of transposition mix (25 μL 2x TD Buffer, 2.5 μL Tn5 transposase (100 nM final), 16.5 μL PBS, 0.1 μL 5% digitonin, 0.5 μL 10% Tween-20, 5 μL H_2_O) and incubated at 37 °C for 30 minutes exactly in a thermomixer with 1000 RPM mixing. DNA was then purified using Zymo DNA Clean & Concentrator-5 Kit to generate 20 μL of product which was pre-amplified for 5 cycles using NEBNext 2x Master Mix (#E7649A) and indexed primers as described previously^44^. A 5 μL of pre-amplified DNA was then used in a 15 μL qPCR reaction using Promega GoTaq qPCR Master Mix to determine the required number of additional cycles to yield minimally amplified DNA for optimal library diversity for next-generation sequencing (determined to be 3-4 additional cycles for these libraries). Amplified DNA was then purified using NucleoMag NGS Clean-up and Size Select beads (Macherey-Nagel) using a 0.5x ratio of beads to DNA to initially clear large DNA fragments from the library and then a 1.5x ratio of beads to supernatant to adsorb DNA fragments of appropriate size for sequencing. DNA-bound beads were then washed twice with 80% EtOH and size-selected DNA was eluted in nuclease-free water. Final libraries were quantified using an Agilent Tapestation using a D5000 screentape and sequenced using an Illumina NextSeq2000 to generate ∼39M paired-end 65bp reads per library.

### RNA-seq

Total RNA was isolated from 1 × 10^6^ cells using RNeasy Plus Mini Kit (Qiagen) with genomic DNA removal. RNA was confirmed to be of high quality using an Agilent Tapestation using an RNA screentape (RIN >9.8 for these samples) and 500ng was prepared for RNA-seq using Truseq RNA library preparation protocol with poly-A capture (Illumina). Final libraries were quantified using an Agilent Tapestation using a D1000 screentape and sequenced using an Illumina NextSeq2000 to generate libraries with 64 – 78 million paired-end 65bp reads.

### Bioinformatics analyses of RNA-seq samples

Alignment of RNA-seq reads to the human reference genome (T2T-CHMv2.0^45^) was performed with ‘subread’ (v.2.0.6^46^) using default options for paired-end read data. Read counting was performed with ‘featureCounts’ with default options for paired-end read data and using strict RefSeq gene annotation. Differential gene expression analysis was performed using the ‘edgeR’ package (v.3.42.4^47^), with lowly expressed genes removed via ‘filterByExpr’ and library normalization with the TMM method^48^. Quasi-likelihood F-tests were performed to assess gene-level differential expression between knock-out and wild type samples while controlling for the false discovery rate at the 0.05 nominal level^49^.

### Bioinformatics analyses of ATAC-seq samples

ATAC-seq sequence reads were trimmed with ‘trimgalorè (v.0.6.10 [https://www.bioinformatics.babraham.ac.uk/projects/trim_galore/]) and aligned to the human reference genome T2T-CHMv2.0 with ‘bowtie2’ (v2.4.4). Post-alignment filtering was performed to remove unmapped reads and read mates, not primary alignments, reads failing platform, orphan reads, read pairs mapping to different chromosomes, and low-quality reads (MAPQ < 30), while keeping only properly paired reads. Duplicated reads were marked and removed with Picard-tools (v.2.26.11 [http://broadinstitute.github.io/picard/]). Peak calling was performed with MACS (v2^50^) in paired-end mode using default options. The number of peaks identified in WT and KO samples was 50,883 and 68,357, respectively. Reads were counted on genome-wide non-overlapping windows of 250bp in size using the ‘csaw’ package (v.1.34.0^51^) and its function ‘windowCounts’ for all human chromosomes. The midpoint of each fragment was used for counting. Below we load the read count data. Background windows were removed via ‘filterByExpr’ and library normalization was performed with the TMM method. Quasi-likelihood F-tests were performed to assess window-level differential accessibility between knock-out and wild type samples while controlling for the false discovery rate at the 0.05 nominal level. Window-level results were summarized to the region-level and promoter-level via ‘mergeResults’ and ‘overlapResults’ functions, respectively. Promoter regions were defined as the 2kbp window around the transcription start site of each annotated gene.

### Bioinformatic analyses of ChIP-seq samples

ChIP-seq sequence reads were trimmed with ‘trimgalorè and aligned to the human reference genome T2T-CHMv2.0 with ‘bowtie2’. Post-alignment filtering was performed to remove unmapped reads and read mates, not primary alignments, reads failing platform, orphan reads, read pairs mapping to different chromosomes, and low-quality reads (MAPQ < 30), while keeping only properly paired reads. Duplicated reads were marked and removed with Picard-tools. Peak calling was performed with MACS (v2^50^) in paired-end mode using default options for MORC2 library, and with options ‘--broad --broad-cutoff 0.1’ for the histone modification H3K9me3 to detect broad peaks. IgG libraries were used as controls during peak calling. The number of peaks identified in H3K9me3 and WT samples were 116,550 and 15,366, respectively.

Data analysis was performed in R (v.4.3.0 [https://www.R-project.org/]). Upset plots were generated with the ‘UpSetR’ package (v.1.4.0^52^). Heatmap and signal profile plots were created with ‘deepTools’ (v.3.5.1^53^) using merged alignments. Read enrichment coverage was computed in 10bp windows using extended and centered reads and using read per genomic content normalization (1x depth).

### Preparation of biotin-labelled λ-DNA

Circular λ-DNA lacking methylation (N3011S, Promega) underwent modification through Taq ligase (M0208L) and New England Biolabs (NEB) buffer (B0208S). This process involved the incorporation of complementary oligomers JT41 (5’(P)-GGGCGGCGACCT-3’(Biotin)) and JT42 (5’(P)-AGGTCGCCGCCC-3’(Biotin)) from Integrated DNA Technologies (IDT). The resultant DNA, now labeled with biotin, underwent purification using the AKTA pure system (Cytiva), followed by size exclusion chromatography using Sephacryl S-500 HR resin (Cytiva).

### Single-molecule measurement and analysis

We adhered to an established protocol for a single-molecule fluorescence assay^18^. In the assay preparation, we created 6 channels on glass slides by drilling 12 holes on the side. The slides underwent cleaning with 10% detergent, acetone, and Mili-Q water, followed by Piranha solution treatment. Subsequently, they were PEGylated using a 1:80 ratio of biotin-PEG to PEG, incubating in a sodium bicarbonate solution (0.1M NaHCO3, pH 8.5) for 12-24 hours. After washing with Milli-Q water and drying with nitrogen gas, additional PEGylation with MS(PEG) solution was performed, with the only difference being the incubation time (1-24 hours). Finally, the channels were partitioned with double-sticky tapes and the edges sealed with epoxy glue.

To immobilize 48.5-kbp double-biotinylated λ-DNA, Streptavidin (100 µg/ml) in T50 buffer (40 mM tris pH7.5 and 50 mM NaCl) was flowed through the channels, incubating for 2 minutes to induce Streptavidin biotin-PEG binding. Channels were washed with T50 buffer, and 48.5-kbp double-biotinylated λ-DNA in imaging buffer (40 mM Tris, 50 mM NaCl, 1 mM MgCl2, 0.5 mM TCEP, BSA (0.25 mg/ml), 50 nM Sytox Orange (SxO, S11368 Thermofisher) was applied at a flow speed of 8 µl/m. After DNA tethering, unbound DNA was removed by flowing imaging buffer with 200 µM Desthiobiotin to dissociate strep-tag of MORC2 from biotin on the surface. For monitoring MORC2-induced DNA compaction, the MORC2 solution (50 nM MORC2 with imaging buffer) was flowed through the channels at a speed of 20 µl/m for 1 minute. Real-time DNA compaction was imaged using a 1mW 561-nm HILO mode laser for SxO excitation. Imaging was conducted with an Olympus UPlanXApo 100x /1.45 lens and an Andor iXon Ultra 897 EM CCD. To obtain kymographs, the intensity values of n pixels (n is proportional to the region of interest (ROI)) from the ROI perpendicular to the extended DNA were summed in each frame. Although the appearance of clusters was sometimes not obvious due to the resolution limit, we determined the compaction time using a fluctuation analysis. Specifically, as the cluster size (or number of clusters) increased, the fluctuation decreased. The term “fluctuation” referred to the variance of the intensity region between adjacent frames.

To assess the kinetics of DNA cluster formation quantitatively, the mean of intensity values below the 50th percentile within the DNA Region of Interest (ROI) was considered as background noise and subtracted from the DNA signal. For a focused examination of DNA variance, the background intensity was standardized to zero (any ‘black (background)’ in the DNA figures signifies zero intensity). To ensure uniform DNA length across all frames, the sum of the DNA signal was equalized after background removal. For calculating the sum of variance intensity, the overlapped region between the current and previous frames was compared, and only the variant segments were utilized.

To determine compaction time, fluctuation data underwent smoothing using Savitzky-Golay with a moving window of 150 or 300 and a polynomial order of 2, followed by normalization. The starting point was established where the DNA signal exhibited a significant initial fluctuation followed by a decrease, indicating the onset of compaction. Finally, an exponential decay equation 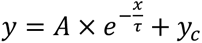 was fitted to the fluctuation data to ascertain the compaction time. If DNA cleavage occurred after complete compaction, *y_c_* was set to zero.

### AFM imaging and analysis

To visualize MORC2/DNA clusters, we combined 10 nM MORC2 with λ-DNA (48.5 kbp from Promega) at a concentration of 10 ng/µL. This mixture was prepared in a reaction buffer (40 mM Tris-pH7.5, 50 mM NaCl, 1 mM MgCl2, 0.5 mM TCEP) in an E-tube and incubated for 10 minutes. The mixtures were then deposited onto poly-L-lysine-treated mica. After a brief wash with 3 mL Milli-Q water, the sample was dried using a nitrogen gun.

AFM measurements of the dried sample were carried out using a Multimode-8 AFM (Bruker) equipped with a Nanoscope VI controller and Nanoscope version 10.0 software. ScanAsyst-Air-HR cantilevers (Bruker, nominal stiffness and tip radius 0.4 N/m, and 2 nm, respectively) were employed. PeakForce Tapping mode was used with an 8 kHz oscillation frequency, and a peak force setpoint value less than 150 pN was applied to minimise sample invasiveness.

For image processing of dry AFM images, Gwyddion version 2.53 was utilized. Initially, background subtraction and transient noise filtering were performed. To ensure exclusive use of the empty surface for background subtraction, masking particles and subtracting background polynomials (planar and/or line-by-line) were implemented, excluding the masked regions. Horizontal scars, arising from feedback instabilities or protein sticking to the AFM tip, were eliminated. Subsequently, plane background subtraction was applied. Finally, blind tip estimation was employed to estimate the tip’s shape, and surface reconstruction was carried out to reduce the broadened effects caused by tip convolution.

For measuring the volume of a single protein, each protein in the images was masked, and grain volume measurement was performed to obtain volume information for each masked protein. For cluster volume measurements, the territory of protein areas bound to DNA was manually defined using Gwyddion. The volume of the defined area was obtained, and to analyse the volume of DNA/MORC2 clusters with a confidence level higher than 99.9%, a threshold of seven standard deviations plus the average of the volume of a single MORC2 was used. Areas with cluster volumes exceeding the threshold were considered clusters.

## Acknowledgements

We acknowledge the use of transmission electron microscopes at the Monash University Ramaciotti Centre for Cryo-Electron Microscopy, and the Ian Holmes Imaging Centre at the Bio21 Molecular Science and Biotechnology Institute. We thank the WEHI Cryo-EM Platform, the WEHI Research Computing Platform and the Monash University MASSIVE high-performance computing facility for providing facilities and support. We acknowledge the WEHI Proteomics facility and Protein Production facility. We would like to thank the Melbourne Mass Spectrometry and Proteomics Facility at The Bio21 Molecular Science and Biotechnology Institute at The University of Melbourne. We acknowledge the technical assistance of Stephen Wilcox, Sarah MacRaild and Lilian Wong. We thank Lori Passmore and MRC Laboratory of Molecular Biology, Cambridge, for purchasing the MORC2 synthetic construct. We thank Matthew Call for comments on the manuscript. We would like to acknowledge Angel Zhuo Chen and Juri Rappsilber for providing suggestions on the quantitation experimental design for crosslinking mass spectrometry and helping set up the data analysis workflow at WEHI. This work was supported by the Charcot-Marie-Tooth Australia Research Grant, and an NHMRC Investigator Grant (2026635) to WT; NHMRC Ideas Grant (2010571) to CRK; FSHD Society Fellowship to AG; NHMRC Investigator Grant (2025645) and Chan Zuckerberg Initiative Grant (2021-237445) to GKS; Snow Medical fellowship and CSL Centenary fellowship to SV; NHMRC Investigator Grant (1194345) to MEB; and NIH NIGMS R35GM133751 grant to SD. We acknowledge the Suh Kyungbae Foundation, Creative-Pioneering Researchers Program through Seoul National University, the Brain Korea 21 Four Project grant funded by the Korean Ministry of Education, Samsung Electronics Co., Ltd. (Project Number IO220811-01964-01), and the National Research Foundation of Korea (Project Number RS-2023-00212694, RS-2023-00265412, RS-2023-00218318) to JKR. SS is supported by funds from WEHI, WEHI Ventures, the estate of Akos and Marjorie Talon, The University of Melbourne Attraction and Retention Funds, the NHMRC Investigator grant (GNT 2016827) and the US Department of Defence Rare Cancer Research Concept Award.

## Author Contributions

WT designed and carried out all biochemical, crosslink mass spectrometry, biophysical experiments, prepared samples for cryoEM and analysed the cryoEM data. HV, AL and SS prepared cryoEM samples and collected cryoEM data. HV and SS analysed cryoEM data. JQL and EH performed the FLIM-FRET and FCS experiments. WT and AG performed the Fluorescence Polarisation ATPase assays. CK and WT performed ChIP-seq, ATAC-seq, RNA-seq data collection. PB, GSK and RSA analysed ChIP-seq, ATAC-seq and RNA-seq data. VV and LD performed proteomics experiment and data analysis. WT, TD and LD performed crosslink mass spectrometry experiments. WT, TD, JY and LD performed crosslink mass spectrometry data analysis. CS and SS performed HDX experiments. CS, SS, PD and SD analysed the HDX data. LC, SS and SV performed MORC2 knocked out experiments. CD provided nucleosomes. MEB contributed to interpretation. JVP, KWM and JKR performed the single molecule DNA compaction assay and atomic force microscopy. SS conceived and supervised the project. WT and SS analysed the data and wrote the manuscript with contributions from all authors.

